# Parallel ecological and evolutionary responses to selection in a natural bacterial community

**DOI:** 10.1101/710715

**Authors:** Elze Hesse, Adela M. Luján, Siobhan O’Brien, Arthur Newbury, Terence McAvoy, Jesica Soria Pascual, Florian Bayer, Dave J. Hodgson, Angus Buckling

**Author notes:** **Corresponding author:** Elze Hesse, CEC & ESI, Faculty of Environment, Science and Economy, University of Exeter, Penryn Campus, Cornwall, TR10 9FE, UK.

## Abstract

Evolution can occur over ecological timescales, suggesting a potentially important role for rapid evolution in shaping community trait distributions. However, evidence of concordant eco-evolutionary dynamics often comes from *in vitro* studies of highly simplified communities, and measures of ecological and evolutionary dynamics are rarely directly comparable. Here, we quantified how ecological species sorting and rapid evolution simultaneously shape community trait distributions by tracking within and between-species changes in a key trait in a complex bacterial community. We focused on the production of siderophores; bacteria use these costly secreted metabolites to scavenge poorly soluble iron and to detoxify environments polluted with toxic non-ferrous metals. We found that responses to copper-imposed selection within and between species were ultimately the same – intermediate siderophore levels were favored – and occurred over similar timescales. Despite being a social trait, this level of siderophore production was selected regardless of whether species evolved in isolation or in a community context. Our study suggests that evolutionary selection can play a pivotal role in shaping community trait distributions within natural, highly complex, bacterial communities. Furthermore, trait evolution may not always be qualitatively affected by interactions with other community members.

**Significance:** Bacterial communities possess remarkable taxonomic and metabolic diversity and play a key role in nearly every biogeochemical process on Earth. Rapid evolution (occurring over ecological time scales) can in principle shape these processes, yet we have little understanding of its importance in natural communities. Here, we quantified how the production of metal-detoxifying siderophores is driven by species compositional changes and evolution in a compost community in response to copper stress. We found that siderophore production converged at intermediate levels, with evolutionary and ecological changes occurring at similar rates. Understanding how ecological and evolutionary processes contribute to shaping trait distributions will improve our ability to predict ecosystem responses to global change, and aid in the engineering of microbial consortia.

## Main text

Changes in community trait distributions are typically driven by inter-specific competition changing the abundance of species carrying certain traits (i.e. ecological species sorting) (1). In the case of microbial communities, which have massive population sizes and short generation times, evolution can occur over timescales commensurate with ecological species sorting (2–4), suggesting a potentially important role of rapid evolution in shaping community trait distributions (5–7) and responses to environmental change (8). Understanding the mechanisms underpinning community trait distributions is not only of fundamental interest, but also has applied relevance (9–11), such as in the engineering of microbial consortia to perform valuable biological tasks (12), including bioremediation (13), and the development of microbiome technologies aimed at improving human health and crop yield (14).

Most work demonstrating a role of rapid evolution in community dynamics has been carried out in *in vitro* environments (2, 15) and/or has relied on the use of greatly simplified communities (3, 16, 17). This may overestimate the role of rapid evolution in community dynamics because, in addition to the typically very strong selection pressures associated with novel *in vitro* environments (4, 18), ecological complexity itself can constrain evolution (19–22) (but see (23)). The response to selection – resulting from ecological species sorting or within-species evolutionary change – typically increases with trait variation (24). Increasing species diversity can both increase between-species trait variation and reduce within-species variation – the latter because of reduced population sizes (25, 26). The relative importance of ecological species sorting versus evolution on shaping trait distributions is therefore likely to increase with species diversity (1, 19). While some recent work has demonstrated rapid evolution occurring over ecological timescales in more complex communities and/or semi-natural environmental contexts (2–4), the role of ecological species sorting versus evolutionary selection in shaping natural community trait distributions is unclear.

A useful model system to track simultaneous ecological and evolutionary changes in functional traits is the ubiquitous production of metal-chelating siderophores in bacterial communities (27). Although perhaps best known for their function as iron carriers (28, 29), siderophores can also protect bacteria from toxic metal stress by preventing the diffusion of toxic metals into bacterial cells (30, 31). Because detoxification takes place in a shared environment, these costly extracellular compounds not only protect the producer, but potentially also neighboring conspecifics (32) and other community members (33, 34). By acting as community-wide public-goods, siderophores play a key role in driving species interactions (35) and adaptive responses to environmental change (34). We previously determined the effect of toxic metals on siderophore production in natural bacterial communities inhabiting mine-degraded soils (33, 36) as well as in experimental compost communities exposed to copper stress (33). In both cases, the presence of toxic metals selectively favored bacterial taxa with greater investment in siderophores, thereby increasing community-wide siderophore levels. However, we have also found that bacterial taxa can benefit from siderophores produced by conspecifics (32) or individuals from phylogenetically diverse taxa (34), suggesting that selection for (higher or lower) siderophore production may also be affected by social interactions.

Here, we investigate ecological changes in siderophore production in response to copper stress in a compost bacterial community simultaneously with evolutionary change in siderophore production by a focal bacterium – the common soil-dwelling bacterium *Pseudomonas fluorescens* (strain SBW25) (37). We hypothesized that ecological changes in siderophore production resulting from exposure to novel copper stress would be greater than evolutionary changes that occurred within SBW25. We also expected the community to alter the evolutionary trajectory of SBW25’s siderophore production as a consequence of both community-imposed evolutionary constraints and greater selection for lower-level siderophore production resulting from the protective effects afforded by other community members.

## Results

### Copper favors siderophore-producing taxa within the compost community

We set up a fully factorial experiment involving the presence/absence of the natural compost community, focal strain *Pseudomonas fluorescens* SBW25 and copper pollution in initially sterile compost (SI Appendix, Fig. S1). These communities were allowed to evolve for six weeks, after which we quantified changes in siderophore production by isolating 24 clones of SBW25 and 24 isolates per replicate community where possible (i.e. *n* = 48 isolates for the SBW25 + community treatments). Isolates were then grown individually in King’s B medium for 48 hrs after which we measured the ability of culture supernatants to chelate iron using colorific CAS assays, corrected for variation in optical density (our measure of siderophore production is based on arbitrary unity; see Methods). We found that copper selected for greater siderophore levels within the compost community compared to unpolluted control communities (Fig. 1A), irrespective of the presence of SBW25 (2-way GLM on mean community siderophore production: *F*_1, 20_ = 0.01, *P* = 0.91 for copper × SBW25, *F*_1, 21_ = 0.04, *P* = 0.85 for SBW25 and *F*_1, 22_ = 8.88, *P* < 0.01 for copper treatment. GLM estimates of mean siderophore production ± 95% confidence intervals [CI] in copper-polluted versus unpolluted communities pooled across SBW25 levels (i.e. presence/absence): 0.55 [0.53, 0.57] and 0.50 [0.48, 0.52], respectively).

**Figure 1.**
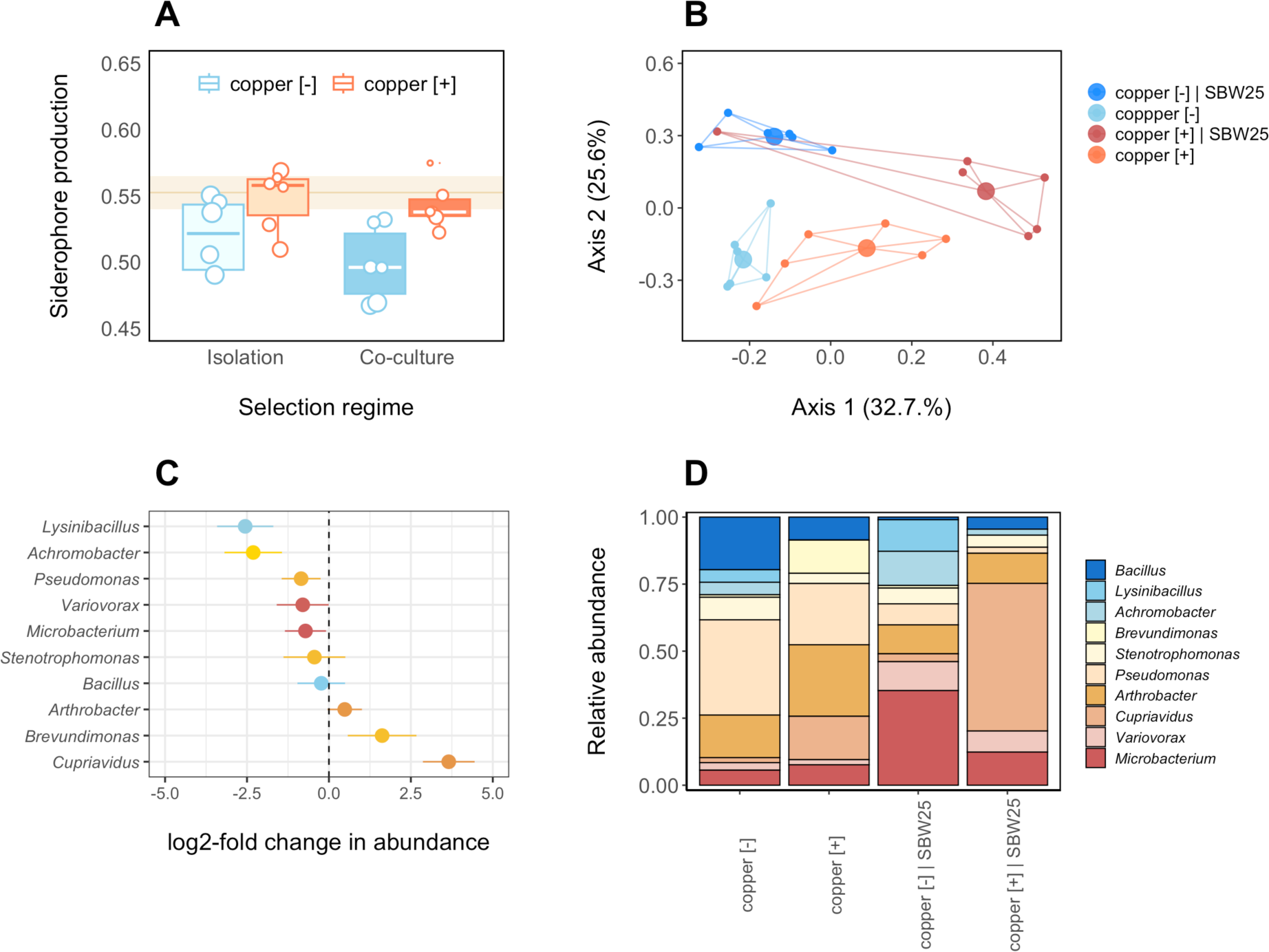
Copper favors siderophore-producing taxa within the compost community. Panel **(A)** shows that copper-polluted (copper [+], orange) communities (*n* =6) produced on average larger amounts of siderophore compared to copper-free (copper [-], blue) communities following six weeks of evolution in isolation (light shading) or in co-culture with SBW25 (dark shading) (*n* = 6 communities per treatment). Brown line and shaded area depict mean ± SE siderophore production of the ancestral communities (*n* = 12) prior to copper amendment. Boxes depict upper and lower quartiles of treatment-specific raw data with the center line showing the median and whiskers providing a measure of 1.5x interquartile range. Points represent the mean value per community based on twenty-four isolates, with within-community variance indicated by different point sizes. Note that for some replicate communities we isolated < 24 isolates, yielding a total sample size of *n* = 548. Panel **(B)** depicts a Principal Coordinate Analysis (PCoA) plot based on Bray-Curtis dissimilarities between compost communities (excluding SWB25). The percentage of variation explained is shown on each axis, calculated from the relevant eigenvalues. Communities belonging to the same treatment are joined with straight colored lines (control_no SBW25_ = light blue, control_SBW25_ = dark blue, copper_no SBW25_ = orange, copper_SBW25_= red), with large points representing treatment-specific centroids and small points individual microcosms. Dissimilarities were calculated using data on the total abundance of different isolates (identified at genus level) within each community that were assayed for siderophore production (*n* = 24 per community). Panel (**C**) depicts the effect of coper on the abundance of bacterial taxa in the compost community. The log2-fold change in total abundance of the ten most common genera when comparing communities that had evolved in unpolluted versus copper-polluted compost. Of the taxa tested, three significantly differed in terms of total abundance between copper-free versus copper-polluted treatments: while *Cupviavidus* was positively affected*, Achromobacter and Lysinibacillus* were both negatively affected by copper stress (see SI Appendix, Table S1 table for Wald test and P-values). Points represent estimated log2-fold change in total abundance of each taxon in response to copper-stress (averaged across culturing conditions) ± standard error bars. Genera are color-coded based on their mean across-treatment siderophore production, ranging from relatively low (blue) to intermediate (orange) and high (red) siderophore production. Panel **(D)** depicts the relative abundance of the ten most common culturable compost taxa, excluding SWB25, following six weeks of evolution under different selection regimes. Genera are listed in order of their across-treatment mean siderophore production, increasing from top to bottom, such that orange-coral taxa produce relatively intermediate to high levels of siderophores.

To determine whether the observed shifts in siderophore production were linked to any compositional changes in the community, we determined the genus-level identity of all final-time-point compost isolates assayed for siderophore production (*n* = 548 isolates in total). We found that copper-mediated changes in siderophore production were accompanied by a shift in community composition (2-way PERMANOVA on Bray–Curtis dissimilarities: copper × social context: *F_1, 20_* = 2.47, *P* = 0.03), with between-community diversity differing significantly across all treatment combinations (pairwise PERMANOVAs: R^2^ > 0.26, all *P*_adj_ < 0.05; Fig. 1B). In particular, the presence of SBW25 changed the relative abundance of taxa within the compost community by suppressing *Pseudomonas, Bacillus* and *Arthrobacter* compared to the community-only treatment (all *P_adj_* <0.05; SI Appendix, Fig. S2). We also found that copper tended to select against taxa that displayed relatively low mean levels siderophore production (i.e. blue taxa in Fig. 1C): in particular, while intermediate siderophore producers, such as *Cupriavidus*, were significantly more abundant in copper-polluted communities, *Achromobacter* and *Lysinibacillus* were negatively affected by copper stress (SI Appendix, Table S1). As a result of such ecological species sorting, the most abundant community members in copper-polluted compost were those displaying more intermediate siderophore levels (Fig 1D). Note that evolution may also have contributed to changes in siderophore production, but this is impossible to quantify from these taxonomic results. We therefore focus on evolution within a focal strain.

### Copper selects against high siderophore levels within SBW25

We tracked siderophore changes within replicate populations of SBW25 that had either evolved in the presence or absence of the compost community. We found the exact opposite response compared to that observed at the community level – toxic copper selectively favored SBW25 individuals that, on average, produced lower amounts of siderophore compared to control populations (LMM: copper effect: 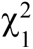 = 10.97, *P* < 0.001. Mean siderophore production ± 95% CI in copper-polluted versus control populations: 0.68 [0.67, 0.70] and 0.72 [0.71, 0.73], respectively). Again, the impact of copper stress on siderophore production did not differ significantly as a function of community presence/absence (LMM: interaction between copper × community: 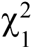 = 0.22, *P* = 0.64; community effect: 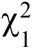 = 1.97, *P* = 0.16; Fig. 2).

**Figure 2.**
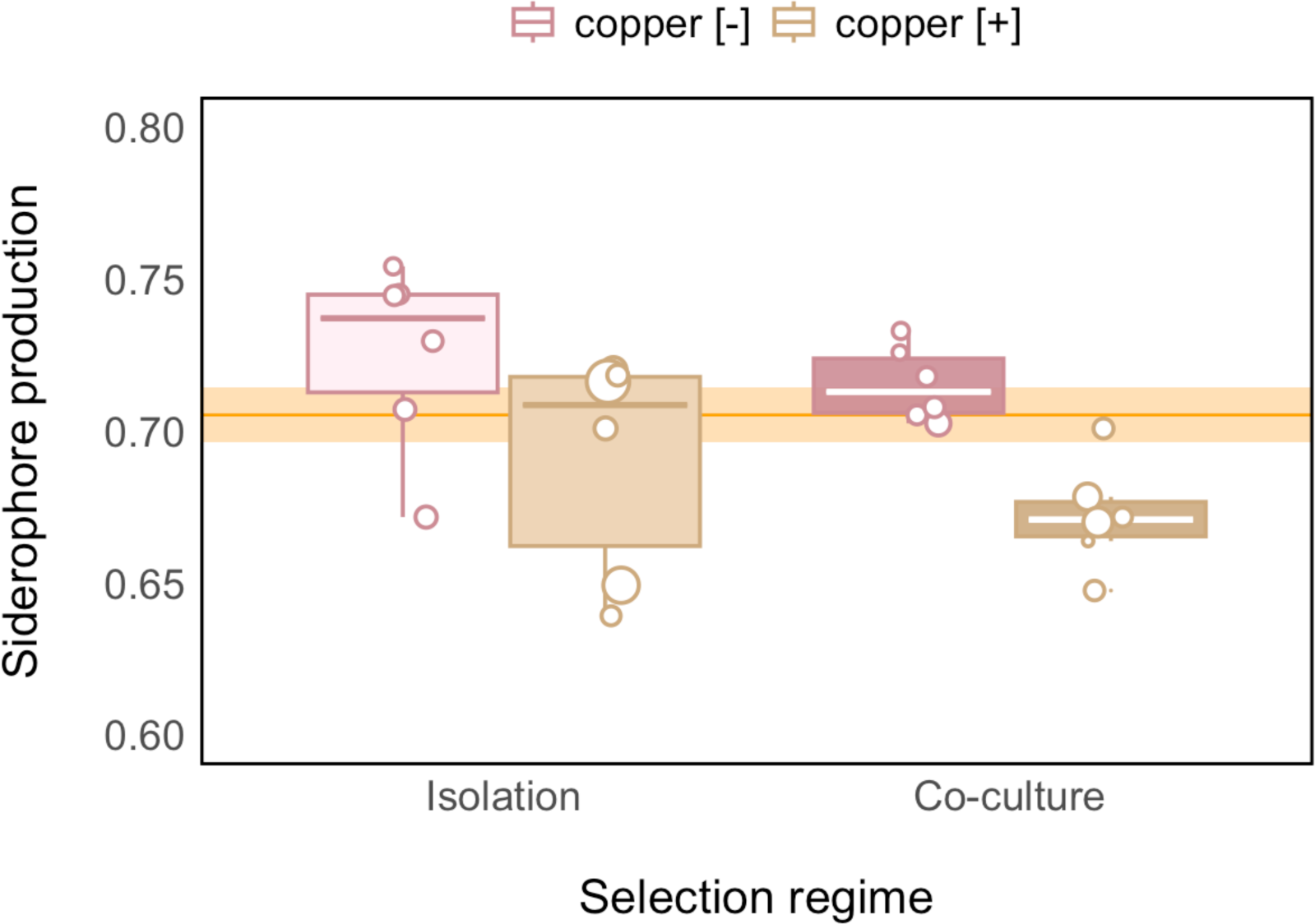
Toxic copper favors the evolution of reduced siderophore production within SBW25. Boxplot showing that copper (copper [+], brown) selected for lower mean siderophore levels in SBW25 populations compared to copper-free conditions (copper [-], pink) irrespective of whether populations evolved in isolation (light-shading) or in co-culture with the compost community (dark-shading) (*n* = 6 populations per treatment). Orange line and shaded area depict mean ± SE siderophore production of ancestral SBW25 populations (*n* = 12) prior to copper amendment. Boxes depict the upper and lower quartiles of treatment-specific raw data with the center line showing the median and whiskers providing a measure of 1.5x interquartile range. Points represent the mean value per population (*n* =6 per copper treatment) based on twenty-four clones where possible, with within-population variance given by different point sizes. Note that for some replicate communities we isolated < 24 clones, yielding a total sample size of *n* = 501.

### Copper selects against siderophore extremes

How can we resolve these apparently opposing responses to copper-imposed selection operating at the species versus community level? To answer this question, we first compared the shape of the distribution of siderophore production of the compost community to that of SBW25. We found that copper selected against siderophore extremes, resulting in trait convergence (Fig. 3), regardless of whether SBW25 and the community were grown in isolation or together in co-culture.

**Figure 3.**
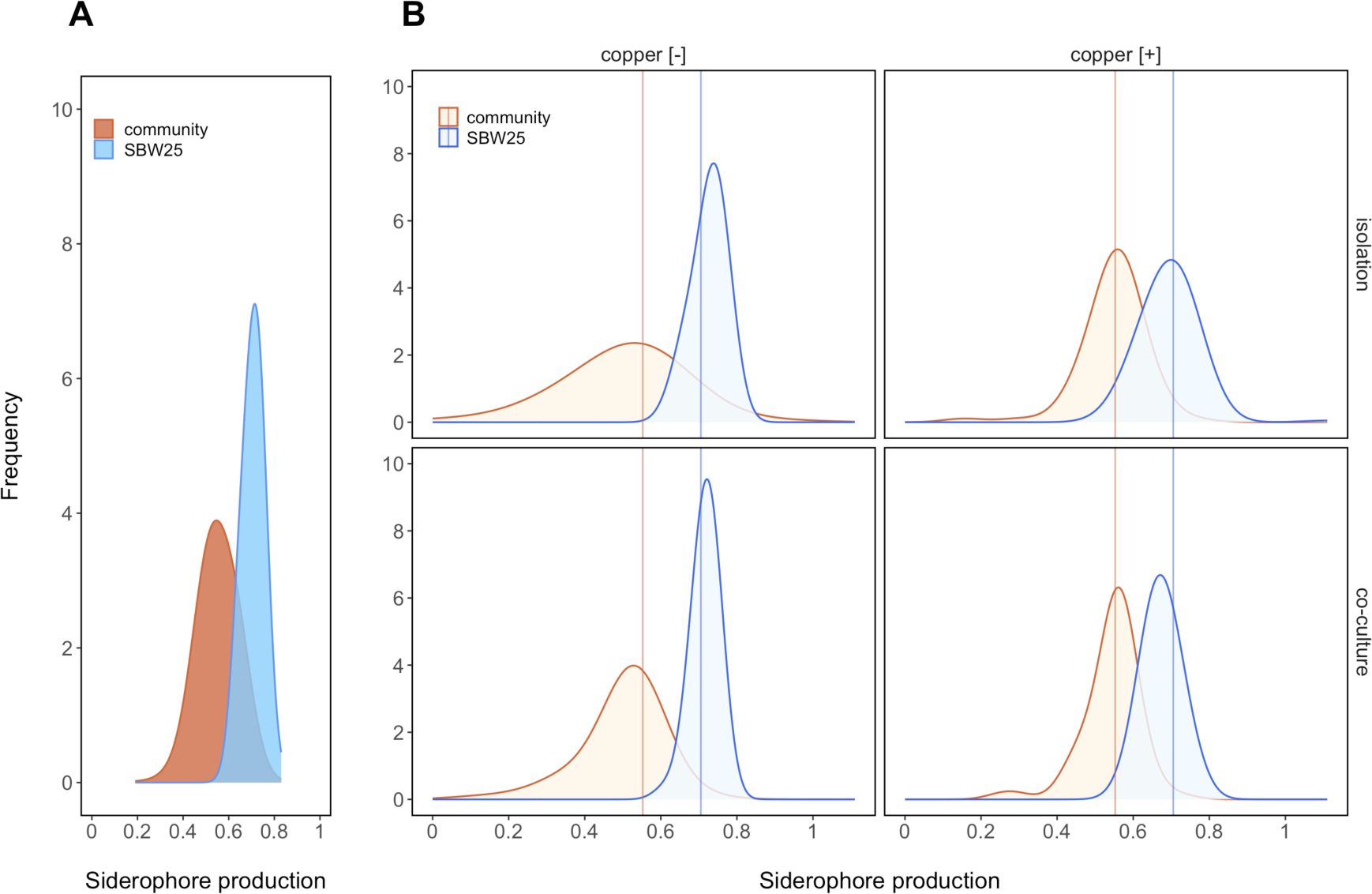
Copper leads to trait convergence by selecting against siderophore extremes. Panel **(A)** shows the frequency distributions of siderophore production in ancestral SBW25 populations (blue) and compost communities (brown) prior to copper amendment. Siderophore data were pooled across copper treatments and replicates (*n* = 24 isolates for each of the twelve replicate SBW25 populations / compost communities). Panel **(B)** demonstrates that the distribution of siderophore production in SBW25 (blue) and the community (brown) converges in response to copper stress, independent of whether these evolved in isolation (top panel) or in co-culture (bottom panel). Siderophore data were pooled across replicates (*n* = 24 isolates for replicate population/community). Note that for some replicates we isolated < 24 clones/isolates of SBW25 or the community, yielding a total sample size of *n* = 1049. Brown and blue vertical lines depict mean siderophore production of the ancestral compost communities and SBW25 populations, respectively.

To quantify this convergence, we focused on the effect that copper has on siderophore production when the community and SBW25 were grown together, using a hierarchical Bayesian linear model to account for heteroscedasticity between isolate types (community versus SBW25; all point estimates are *maximum a posteriori estimates* ± 95% credible intervals). The copper-mediated change in siderophore production in SBW25 was −0.042 [-0.072, −0.014]. In contrast, copper stress increased siderophore production among compost community isolates by 0.044 [0.009, 0.081]. Thus, the effect was of approximately equal magnitude, but in the opposite direction. Indeed, combining the posterior estimates for the effect of copper on siderophore production in SBW25 and in the rest of the community (positive values indicate that the positive effect on community isolates is larger than the negative effect in SBW25 and *vice versa*), the estimated combined effect was 0.001 [-0.044, 0.045]. In the absence of copper, the difference in siderophore production between SBW25 and the compost community was 0.216 [0.185, 0.251]. Copper pollution significantly reduced this figure by 0.086 [0.041, 0.133] to 0.130 [0.098, 0.161]. Thus, opposing and approximately equal effects of selection lead to a significant reduction in the difference between (relatively high producing) SBW25 and (relatively low producing) community isolates under copper stress.

The model described above included separate standard deviation parameters for each sample type (community/SBW25 × copper/no copper) to estimate their respective within-microcosm variation in siderophore production. The standard deviation of siderophore production amongst SBW25 isolates was 0.0294 [0.0267, 0.0340] in the absence of copper. The copper treatment led to an increase of 0.006 [-0.00031, 0.0131] to 0.0354 [0.0315, 0.0427]. Conversely, variation in siderophore production was reduced amongst compost community isolates in the presence of copper. Here, the standard deviation fell from 0.0994 [0.0899, 0.115] to 0.0693 [0.0633, 0.0808] in response to the copper treatment, a significant difference of 0.0301 [0.0157, 0.0474]. That the overall variation in siderophore production was much higher in the compost community than SBW25 is expected, since measurements were taken from a range of unrelated strains. The decrease in variance due to copper stress suggests ecological species sorting (and some evolutionary change), whereas the increase in variance within SBW25 isolates is compatible with evolutionary change.

To further investigate the generality of this result, we focused on isolates of representative genera (*n* = 8 genera) that occurred in both copper environments and had been assayed for siderophore production. We found that trait changes were qualitatively similar: copper selected against siderophore extremes. That is, taxa with high ‘baseline’ siderophore levels (i.e. mean siderophore levels > 0.55 in control conditions) produced fewer siderophores in copper-polluted compost (Cohen’s *d* < 0; SI Appendix, Fig. S3). In contrast, copper stress favored greater siderophore production in bacterial taxa with low ‘baseline’ siderophore levels (Cohen’s *d* > 0). Hence, we found a negative relationship between effect size and ‘baseline’ siderophore levels (Spearman rank correlation: *S* = 624.36, *P* = 0.005, *ρ* = - 0.72).

### Direct fitness costs of high levels of siderophore production in copper

Selection for siderophore production in the presence of toxic copper can be explained by the direct protective effects afforded by siderophores. However, what is less clear is why there was selection against *high* levels of siderophore production in toxic copper conditions. There are two general, non-mutually exclusive explanations. First, there may be a direct cost associated with greater investment in siderophores in copper when individuals produce more than enough siderophore to detoxify copper, making the excess an unnecessary metabolic burden. Second, individuals producing high levels of siderophore may protect competing conspecific and heterospecific individuals, increasing the relative cost of siderophore production.

We first explored how evolved changes in siderophore production affect individual performance in toxic copper conditions. Specifically, we determined whether SBW25 clones that had evolved to produce relatively low levels of siderophore displayed lower copper tolerance (*n* = 277 clones in total, all isolated from the ‘no community’ treatment; SI Appendix, Fig S1). To this end, evolved clones were individually grown in King’s B broth supplemented with a toxic dose of copper sulphate (final concentration of 6.17 mM CuSO_4_). We recorded changes in optical density (OD_600_) over a 48-hrs period and estimated clone-specific Malthusian growth parameters (*m*). Our results show that clones that had evolved in copper-free and copper-polluted compost grew equally well in toxic copper broth (LMM: effect of evolutionary background on *m*: 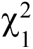 = 0.76, *P* = 0.39; SI Appendix, Fig. S4), suggesting that siderophores are effective in detoxifying copper. Furthermore, reduced investment in siderophores did not confer lower tolerance, irrespective of a clones’ evolutionary history (LMM on *m*: siderophore production × copper background: 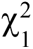 = 0.02, *P* = 0.88; copper background: 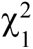 = 0.16, *P* = 0.69; siderophore production: 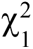 = 2.95, *P* = 0.09; SI Appendix, Fig. S4). The observed shift towards reduced siderophore levels in SWB25 therefore did not alter copper tolerance.

We next investigated why high investment in siderophores is costly in the presence of copper but not in the absence. We speculated that the constrained growth of SBW25 in the presence of copper (SI Appendix, Fig. S5) may have reduced levels of siderophore required for the canonical function of siderophores: iron uptake (38). To test this hypothesis, we manipulated population growth rates (and hence indirectly the need for iron to sustain growth) by inoculating monocultures of the wild-type pyoverdine-producing SBW25 strain and an isogenic pyoverdine-deficient mutant (PBR840) (39) in sterile compost at very low density (allowing for faster growth) and high density (allowing for slower growth), both in the presence and absence of copper pollution. This mutant does not produce the main siderophore pyoverdine (which reduces total siderophore production to 21% of the wild-type) and displays levels of siderophore production similar to an average compost community member.

Confirming our observations on copper tolerance in evolved SBW25 populations, the pyoverdine-deficient mutant performed equally well compared to the pyoverdine producing strain in copper-polluted compost (GLM on *m*: copper × strain: *F_1,43_* = 0.52, *P* = 0.48; Fig. 4A), with copper significantly reducing population growth (main effect copper: *F_1,44_* = 100.24, *P* < 0.001. Mean growth ± 95% CI: 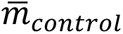 = 1.50 [1.45, 1.55] and 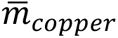 1.14 [1.09, 1.19], pairwise contrast: *t* ratio = 10.01, *P* < 0.001), irrespective of inoculation density (copper × density: *F_1,40_* = 0.22, *P* = 0.65). We also found that in both copper environments, the pyoverdine producers benefitted when inoculated at low density and hence growth was not constrained (strain × density: *F_1,43_* = 16.30, *P* < 0.001; 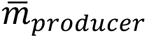 = 1.98 [1.90, 2.05] and 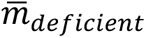 1.88 [1.81, 1.95], pairwise contrast: *t* ratio = 1.89, *P* = 0.07; Fig. 4A). In contrast, the non-producing mutant grew faster than the producer under slow (high inoculation density) growth conditions (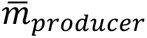 = 0.61 [0.54, 0.69] and 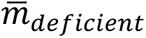 = 0.81 [0.74, 0.88], pairwise contrast: *t* ratio = −3.82, *P* <0.001). Consistent with this monoculture pattern, the relative fitness of the pyoverdine producer was greater when competing with the mutant under high growth conditions (low inoculation density), while it was lowest under low growth conditions (GLM on 𝑟*_producer_* : density × copper interaction: *F_1,19_* = 6.95, *P* = 0.016) – especially in the presence of copper (high density: 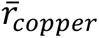 = 0.79 [0.73, 0.85] and 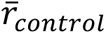 = 0.91 [0.85, 0.97], pairwise contrast: *t* ratio = 2.92, *P* < 0.01 and low density: 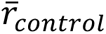 = 1.04 [0.98, 1.10] and 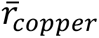 = 1.00 [0.94, 1.07], pairwise contrast: *t* ratio = −0.86, *P* = 0.40. Data on Malthusian growth and relative fitness are presented in SI Appendix, Fig. S6). These results suggest a cost to high investment in siderophores when growth is otherwise constrained, while there is no cost and potentially a benefit to high levels of siderophore production when growth is less constrained, and these compounds are needed to compete for iron.

**Figure 4.**
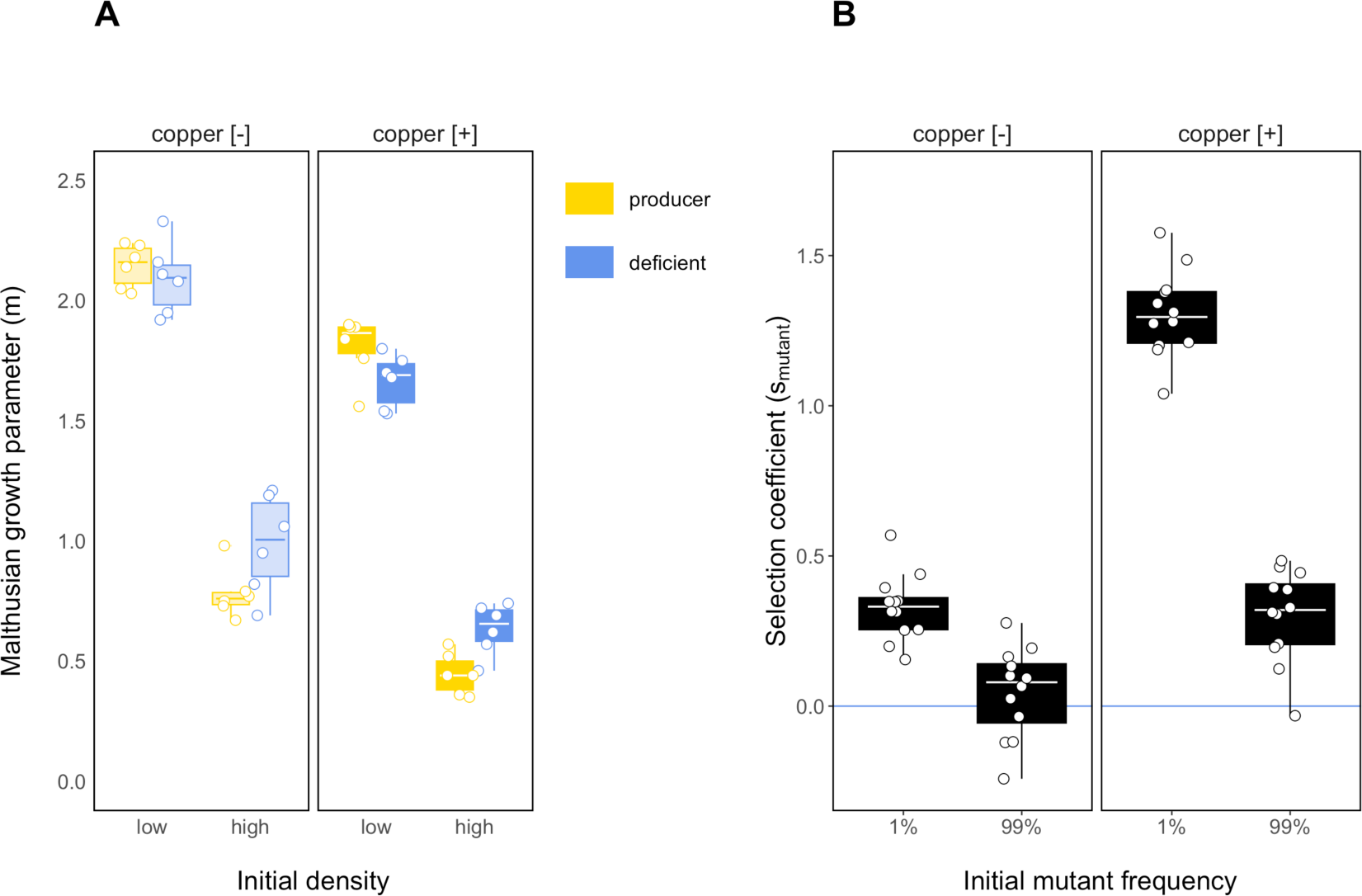
Benefit of different siderophore strategies is context dependent. Boxplot showing **(A)** the Malthusian growth parameter (*m*) of the pyoverdine-deficient mutant (blue) and pyoverdine-producing wild-type SBW25 strain (gold) as a function of inoculation density when individually grown in copper-polluted (copper [+]) and unpolluted (copper [-]) compost (**B)** the pyoverdine-deficient ‘cheat’ receiving a relative selective benefit (𝑆_𝑚𝑢𝑡𝑎𝑛𝑡𝑠_ > 0) when rare, in particular when competing with the wild-type producer in copper-polluted compost. Strains were co-inoculated together at different frequencies (1% or 99%) in copper-polluted (copper [+]) and unpolluted (copper [-]) compost. Blue horizontal line depicts equal fitness (𝑆_𝑚𝑢𝑡𝑎𝑛𝑡𝑠_ = 0) of competing strains. Boxes depict the upper and lower quartiles of treatment-specific raw data with the center line showing the median and whiskers providing a measure of 1.5x interquartile range, and points show the raw data for each replicate population.

### Public-goods exploitation favors reduced investment in siderophores

While direct costs and benefits could qualitatively explain our results, we next explored whether public-goods exploitation could have played an additional role in driving selection against SBW25 individuals producing relatively large amounts of siderophores. Previous work investigating within-species exploitation of siderophores has demonstrated that siderophore production is costly to the individual (40, 41), creating incentives for non-producing ‘cheats’ to invade and steal these public-goods away from the producer (42). We have also found that bacterial compost isolates that invest relatively little in siderophores benefit from other species’ detoxifying siderophores under similar levels of copper stress (34). However, siderophore-mediated facilitation could not fully rescue the growth of copper-sensitive isolates, nor did it result in exploitation (i.e. no detectable indirect costs by helping competitors) within the community over ecological timescales (34). Combined with our current finding of parallel changes in siderophore distribution at the within-taxon and community level (Fig. 3), we focus on social dynamics *within* species only.

To determine whether individuals with lower siderophore investment can exploit relatively high producers we carried out competition assays. In particular, we manipulated the relative frequencies of the wildtype and pyoverdine-deficient mutant of SBW25 (1% or 99% starting frequency of strains) and assayed their relative performance following 7 days of growth in copper-polluted and unpolluted compost. Negative frequency-dependent fitness often characterizes social interactions in structured environments, where rare non-producing ‘cheats’ are more likely to readily encounter cooperators rather than other cheats and grow relatively well, while rare cooperators will mostly encounter cheats, but so will other cheats; with the latter suffering a greater relative growth cost. We found that the pyoverdine-deficient mutant experienced a relative selective benefit when rare, in particular in toxic copper conditions (GLM on selection coefficient (*s*) of pyoverdine-deficient mutant: frequency × copper: *F*_1, 44_ = 79.30, *P* <0.001. Mean ± [95% CI] for control: 𝑠_*high*_= 0.045 [-0.08, 0.13] and 𝑠_*low*_ = 0.33 [0.25, 0.41]; pairwise contrast; *t* ratio = −4.95, *P* < 0.001. Mean ± [95% CI] for copper: 𝑠_*high*_= 0.301 [0.22, 0.38] and 𝑠_*low*_ = 1.31 [1.22, 1.39]; pairwise contrast; *t* ratio = −17.55, P < 0.001; Fig. 4B. Data on Malthusian growth are presented in SI Appendix, Fig. S7). This shows that in addition to direct fitness effects, social exploitation by conspecifics could also have contributed to the evolution of reduced siderophore production by SBW25 in response to copper stress.

## Discussion

Using siderophores as a focal trait, we show that evolutionary and ecological processes operate in parallel and over similar timescales in a natural bacterial community, resulting in trait convergence towards intermediate siderophore levels in response to copper stress. In particular, copper-imposed selection changed the relative abundance of bacterial taxa by selecting against individuals that displayed extremes in siderophore investment. This pattern was consistent at the community- and within-taxon level. Crucially, changes in siderophore production not only resulted from ecological species sorting, but also from evolutionary changes within species: SBW25 adapted to copper stress by lowering its investment in siderophores to levels more comparable to that of an average compost community member.

In our system, copper pollution resulted in selection against siderophore extremes. We propose such stabilizing selection in part resulted from directly incurred costs borne by taxa with relatively low and high siderophore production. In particular, we have previously shown that bacterial compost taxa that invest relatively little in siderophores display reduced copper tolerance and suffer an absolute growth cost (34). Our current findings indicate that beyond a certain threshold, increased investment in detoxifying siderophores resulted in diminishing returns, thereby selecting against individuals producing relatively large amounts of siderophore.

This raises the question as to why siderophore production was maintained at higher levels in unpolluted compared to copper-polluted environments. It has been well-established that siderophores play an important role in scavenging insoluble iron from the environment (43). Ironbound siderophores are typically imported by specific receptors that constrain their use by other competing strains, thereby limiting siderophore cross-feeding between strains (but see (44, 45)). As such, siderophores are thought to play a key role in determining the outcome of interspecific competition for iron (46, 47). Hence, siderophores were likely positively selected for in copper-free conditions to pillage iron from competing strains. By contrast, competition for iron was likely reduced under copper stress because iron no longer limited bacterial growth rates (copper did); the main role of siderophores in copper-polluted conditions was to detoxify metals.

Social conflict and cooperation are often thought to play a key role in driving the eco-evolutionary dynamics of microbial communities (48–53). In microbes, social behaviors are often mediated by secretion of ‘public-goods’ – metabolically costly compounds that benefit the producer, but that are open to exploitation by non-producers that do not contribute their fair share but utilize the benefits afforded by their production (54). Most of our understanding of public-good dynamics comes from work on within-species interactions (55), yet many public-goods can simultaneously benefit other species (56–60), particularly in microbial communities. Key examples include antibiotic-degrading enzymes (61, 62), resource-scavenging molecules (55, 63), immune-manipulating effectors (64–67) and potentially siderophore-mediated detoxification. However, we found little evidence to suggest that inter-specific facilitation and exploitation shaped siderophore production in this experiment: siderophore production of neither the community nor SBW25 was qualitatively affected by the other, despite – and in line with previous work – SBW25 greatly altering community composition over the course of our experiment (68). We did find some evidence for within-species exploitation: the pyoverdine-deficient ‘cheat’ accrued a relative fitness benefit when competing with wild-type pyoverdine-producing SBW25, in particular when rare, both in the presence and absence of copper. Such frequency dependent dynamics are indicative of intraspecific exploitation of public-goods in spatially structured environments (55, 69–71), and is predicted to increase variation within species while decreasing divergence between species, which is borne out by our results.

In our experiment we quantified the *potential* of bacterial isolates to produce siderophores rather than their *in-situ* production in compost. The transcription of genes encoding the proteins that participate in siderophore biosynthesis is tightly regulated by the presence of iron and/or toxic metals (30). However, the use of a common garden environment to quantify siderophores was necessary because: (1) for the majority of environmental isolates we do not exactly know what kind of metal-chelating agents they produce, meaning a metagenomics approach is currently not viable; (2) it is notoriously difficult to extract siderophores from soil and link their production to specific taxa and (3) the presence of soil organic material and other factors in compost interferes with CAS assays. Notwithstanding, we demonstrate a direct link between relative changes in siderophore potential and fitness.

Our focus on public-goods traits, such as siderophore production, may have over-estimated the role of evolution. Previous work has shown that public-goods cheats readily emerge via loss-of-function mutations, including cheats defective for quorum sensing (72) or the production of extracellular proteases (73), polysaccharides (74) and siderophores (70). For example, pyoverdine-deficient cheats typically evolve within days in the lab (75), often via single SNPs in regulatory genes. However, over-estimation of the role of evolution may be mitigated to some extent because, compared to many studies on the evolution of public-goods, we evolved compost communities over an extended period which also allowed for major shifts in community composition. More broadly, adaptive evolution of a wide range of non-social traits can result from loss-of-function mutations through rewiring of regulatory and metabolic networks (76). Finally, gain of functions, including potential public-goods traits, can also potentially evolve very rapidly via horizontal gene transfer (HGT; 77, 78, 79) (80). HGT can occur across large phylogenetic distances (81), which could actually increase the relative importance of evolution *versus* species sorting in shaping community trait distributions.

To conclude, we show that the importance of rapid evolution in shaping community trait distributions is not limited to simplified *in vitro* communities but can also play a role as important as species sorting within highly complex natural microbial communities. Furthermore, our results suggest that trait evolution may not always be qualitatively affected by interactions with other community members.

## Methods

All supporting data and R code are available at https://zenodo.org/records/11639136. Sequences have been deposited at ENA Project PRJEB29924: https://www.ebi.ac.uk/ena/data/search?query=PRJEB29924.

### Bacterial strains

We used *Pseudomonas fluorescens* strain SBW25 as our focal species (37). This common soil-dwelling bacterium produces a variety of siderophores, including pyoverdine (45) and an ornicorrugatin-like siderophore (82), that are able to chelate iron. The strain’s primary siderophore – pyoverdine – has been shown to bind copper (30), which renders the environment less toxic by preventing diffusion into the bacterial cell (31). To facilitate isolation of SBW25 from the community, the strain was modified by inserting a LacZ genetic marker and a gentamicin-resistance cassette, which resulted in blue resistant colonies when plated out on Lysogeny Broth (LB) agar supplemented with 90μg/mL 5-Bromo-4-chloro-3-indolyl-β-D-galactopyranoside (X-gal) and 25 μg/mL gentamicin (32).

To isolate the compost community, 40 gr of fresh compost (Verve John Innes No. 1) was added to 200 mL of M9 minimal salt solution and incubated shaken at 150 rpm at 28 °C for 24h. The supernatant was plated out on LB agar to verify the presence of a bacterial community, and also on LB supplemented with 30μg/mL gentamicin to verify the susceptibility of the community to this antibiotic (32). A 20% glycerol stock solution was prepared from the aqueous sample suspension and frozen at −80°C for future use.

### Selection experiment

Compost microcosms were established with 30 gr of twice-autoclaved sterile compost in round 90 mm petri dishes. To track ecological and evolutionary changes in siderophore production, we seeded twelve replicate compost microcosms with (1) a soil wash of the isolated compost community, (2) an overnight culture of SBW25 or (3) both together at a 1:1 ratio (*n* = 36 compost microcosms in total, SI Appendix, Fig. S1), keeping total inoculation density constant (∼5 × 10^7^ cells). Note that results of the community-only treatment were published previously (33). Compost microcosms were incubated at 26°C and 75% relative humidity for 24h, after which half of the microcosms of the above treatments (*n* = 6) was supplemented with a toxic dose of copper (2 mL of filter-sterilized 0.25M CuSO_4_) and the remainder with an equal volume of sterile *dd*H_2_0 (^3^_3_). Microcosms were incubated for a total of 6 weeks. After three weeks, a second dose of copper or *dd*H_2_O was added to the relevant microcosms. Samples were taken just before (‘ancestral’) and 6 weeks after copper amendment (‘evolved’) by transferring 1 gr of compost per microcosm to 6 mL of M9 solution in 30 mL glass vials. Vials were shaken for 2h at 26°C at 180 rpm, after which supernatants were frozen in glycerol (25% final concentration) at - 80°C for future assays.

### Phenotypic assays on ancestral and evolved isolates

#### Siderophore production

To quantify siderophore production, serial-diluted freezer stocks (taken just before and 6 weeks after giving the first dose of copper) were plated out on duplicate LB agar plates supplemented with (i) X-gal to isolate members of the compost community or (ii) X-gal and gentamicin to isolate SBW25 colonies. Following 48 hours of incubation at 26°C, individual colonies could be identified, counted and picked. For each unique treatment-time combination, we picked 24 colonies of the compost community and/or SBW25 per replicate, with a total of 48 colonies picked from compost microcosms seeded with both SBW25 and the community. Individual colonies were grown in KB broth for 48h at 26°C, after which the supernatant was assayed for the extent of iron chelation. Siderophore production was quantified using the liquid CAS assay described by Schwyn and Neilands (^8^_3_). Siderophore production per isolate was estimated using: [1 − (A_i_/A_ref_)] /(OD_i_), where OD_i_ = optical density at 600 nanometer (nm) and A_i_ = absorbance at 630 nm of the assay mixture and A_ref_ = absorbance at 630 nm of reference mixture (KB+CAS; A_ref_). For some samples, absorbance reads were negative following reference correction, so we standardized siderophore production by setting the minimum observed value to zero. Note that CAS assays performed in iron-limited KB medium provided qualitatively similar results (34).

#### Copper tolerance

We determined the direct fitness consequences of copper-imposed reductions in siderophore production in SBW25. Final-time-point clones (*n* = 277) isolated from populations that had evolved in the absence of the community were grown individually at 26°C for 24h, after which ∼ 10^4^ cells were inoculated into 96-well plate wells containing 200 μL of King’s B broth supplemented with a toxic dose of copper sulphate (final concentration of 6.17 mM CuSO_4_). Individual clones were incubated statically at 26°C for 52 hours, and their optical densities (OD_600_) were measured every 8–12 h (Varioskan Flash plate reader, Thermo Scientific, Waltham, MA, USA). We determined the Malthusian growth parameter (*m*) for each clone as: ln(final density / start density) / time (hours) (84).

### Sanger sequencing of compost isolates

The 16S rRNA gene of all final-time-point compost isolates assayed for siderophore production was sequenced to confirm genus-level identity (i.e. *n* = 24 per replicate). PCRs were performed in 25µL reactions containing 1x DreamTaq Green PCR Master Mix (2X) (Thermo Scientific), 200 nM of the 27F and 1492R primers and 3 µL of 1:100 diluted culture that had undergone 3 freeze-thaw cycles. The thermal cycling parameters were set to 94°C for 4 min, followed by 35 cycles of 1 min at 94°C, 30s at 48°C and 2 min at 72°C, and a final extension of 8 min at 72°C. Following Exo-AP clean-up, high quality samples were Sanger sequenced using the 27F primer (Core Genomic Facility, University of Sheffield). Sequence quality was assessed using the R ‘dada2’ package (85), and sequences were trimmed in Genious (86) to achieve an overall quality score >35. Using Mothur (87), sequences longer then 300bp were then aligned to the Silva.Bacteria.Fasta database, and taxonomy was classified using the RDP trainset 14 032015 as a reference database.

### The role of siderophores as iron-scavengers as a function of density

To determine whether high siderophore production was selected against as a result of iron being less limited under toxic copper conditions, we conducted competition assays between the pyoverdine-producing LacZ-marked SBW25 strain (37) and an isogenic pyoverdine-deficient mutant (PBR840) (39) in 30 mL glass universals containing 5 grams of twice-autoclaved sterile compost. Using a full-factorial design, strains were grown in isolation or together at a 1:1 ratio at either low (10^2^ CFUs) or high inoculation density (10^6^ CFUs), keeping total inoculation density constant across these social contexts. Compost microcosms were incubated for 24 hours at 26°C and 75% relative humidity, after which we added a toxic dose of CuSO_4_ (1 mL of 0.25M CuSO_4_) to half of the microcosms and an equal volume of sterile *dd*H_2_O to the remainder *(n* = 6 replicates per treatment). Compost microcosms were incubated for a total of 7 days, after which soil washes were taken, serial dilutions of which were plated out on KB agar. For co-cultures, strains could be distinguished based on their colony pigmentation: wild-type colonies were blue on X-gal supplemented KB agar, whereas mutant colonies were white. We determined the Malthusian growth parameter (*m*) for each strain as described above.

### Public-goods exploitation in SBW25

To determine whether social exploitation could have played an additional role in driving copper-mediated reductions in siderophore production in SBW25, we conducted competition assays between the pyoverdine-producing wild-type and pyoverdine-deficient mutant strain of SBW25 in petri dishes containing 30 gr of twice-autoclaved sterile compost. Using a full-factorial design, strains were inoculated together at a low (1%, 10^4^ CFUs, *n* = 24) or high (99% 10^6^ CFUs, *n* = 24) frequency. Compost microcosms were incubated for 24 hours at 26°C and 75% relative humidity, after which we added a toxic dose of CuSO_4_ (2 mL of 0.25M CuSO_4_) to half of the microcosms and an equal volume of sterile *dd*H_2_O to the remainder. Following 7 days of incubation, soil washes were taken and plated out on KB agar supplemented with X-gal to distinguish between strains. We determined the Malthusian growth parameter (*m*) for each strain as above.

### Data analyses

For all analyses, we used *R* Version 4.0.3 (R Development Core Team; http://www.r-project.org). In general, models were compared by sequentially deleting terms and comparing model fits using F-tests or χ^2^-tests (where appropriate), after which pairwise contrasts were computed using the *‘*emmeans’ packages (88), with α < 0.05. In case of multiple pairwise testing, *P*-values were adjusted using the false discover rate (‘fdr’) method unless stated otherwise. We checked residual behaviour using the ‘DHARMa’ package (89). All plots were produced using the ‘ggplot2’ package (90).

#### The effect of copper on community structure and mean levels of siderophore production

To test for the effect of copper on siderophore production in compost communities that had either evolved with or without SBW25, we initially used a linear mixed effects (LMM) model (*‘lmer’* function in *‘lme4’* package) (91) with copper × SBW25 presence as fixed explanatory variables, as well as their interaction. To account for non-independency of observations we fitted random intercepts for individual microcosms. However, we observed strong heteroscedasticity of residuals, which could not be remedied by including a treatment-specific variance structure. For each replicate community, we therefore averaged siderophore production across the 24 compost isolates assayed and used a 2-way generalized linear model (GLM) with a Gaussian error distribution to test for the interactive effect of copper × SBW25 presence on mean community-wide siderophore production. This simplified model gave qualitatively very similar results to the LMM without violating any underlying model assumptions.

We determined the genus-levels identity of all end-point compost isolates assayed for siderophore production (*n* = 24 per community) and used PCoA ordination plots to depict pair-wise Bray-Curtis dissimilarities in community composition between microcosms (excluding observations on SBW25). To test for the interactive effect of copper × SBW25 presence on community structure we ran a permutational 2-way ANOVA test on Bray-Curtis dissimilarities using the ‘*adonis2*’ function in the ‘*vegan*’ package (92) with 9999 permutations. Following this, we ran pairwise permutational ANOVAs to disentangle which treatments differed significantly from one another using the *‘calc_pairwise_permanovas’* function in the ‘*mctoolsr’* package (https://github.com/leffj/mctoolsr/), with *P-*values corrected for multiple testing using the ‘fdr’ method. We confirmed homogeneity of variance using the *‘betadisper’* function in the ‘*vegan’* package to estimate group differences in dispersion and treated all copper × SBW25 combinations as levels of a single factor in a 1-way AVOVA on dispersion.

To determine which of the compost isolates differed in abundance across treatments, we fitted a negative binomial GLM to the data using the ‘*DESeq’* function in the R package ‘*DESeq2*’ (93). We focused on a subset of ten common compost isolates that occurred in both unpolluted and copper-polluted communities. Based on the outcome of pairwise permutational ANOVAs (see above), we initially included the interaction term between copper × SBW25 presence in the design of our GLM. However, following model reduction (LRT tests), the minimal adequate model was best described by only including the additive effect of copper and community context. Using this additive model, we calculated significant differences in the abundance of taxa using Wald tests and we corrected *P* –values for multiple testing for each of two contrasts used (presence/absence of either SBW25 or copper).

#### The effect of copper on mean siderophore production in a focal strain

To test for the interactive effect of copper and community context on siderophore production in SBW25, we used a LMM model with copper × community presence as fixed explanatory variables, as well as their interaction. To account for non-independency of observations, we fitted random intercepts for individual microcosms. Based on the obtained simulation-based residual plots, we included a dispersion parameter for each of the copper × community levels, using the *‘glmmTMB’* function in the *‘glmmTMB’* package (94).

#### The effect of copper on the distribution of siderophore production

To demonstrate that copper selectively favored intermediate siderophore levels, we determined the effect size of copper for bacterial taxa that commonly occurred in both copper treatments and related this to their mean siderophore production under non-polluted conditions, using a Spearman rank correlation. Effect size was calculated using the *‘cohen.d’* function in the *‘effsize’* package (95). Cohen’s *d* takes the difference between two means and expresses it in standard deviation units, with negative values indicating a reduction in siderophore levels in response to copper, and *vice versa* for positive values.

#### Effect-size in compost communities evolved with SBW25

To determine the impact of copper on the siderophore production of both SBW25 and community isolates that had evolved together, we used a hierarchical Bayesian linear model which accounts for the differing variances associated with explanatory variables (i.e. siderophore measurements from community isolates were taken from diverse bacterial strains even within a single replicate community, as opposed to the originally isogenic SBW25 stocks):

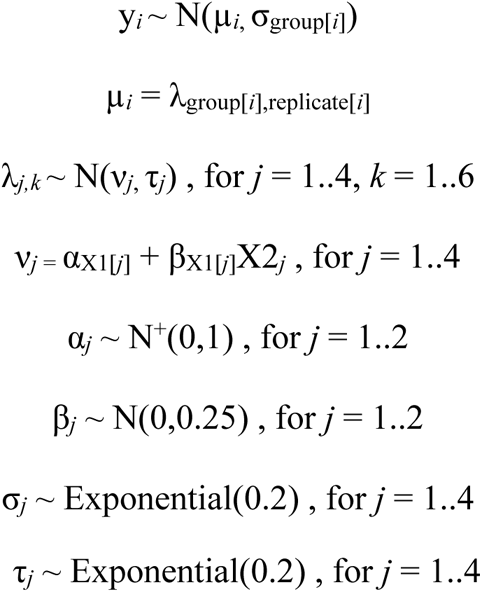

where X1 is a vector of indices indicating SBW25 vs community samples; X2 is a vector indicator indicating the presence/absence of copper; *replicate* and *group* are vectors of indices denoting the replicate microcosm (from a given group) and the combination of isolate type (SBW25 vs community) and presence/absence of copper for each data point. The model above, with separate standard deviation terms (σ) for each group was chosen over models with one or two (determined by isolate type or copper presence) standard deviation terms after model selection by Pareto smoothed importance sampling leave one out (PSIS-LOO) cross validation (96). Inference was performed by Markov chain Monte Carlo using Turing.jl (97) in the Julia programming language (98): 4 chains of 1000 samples each using the No U-Turn Sampler (NUTS) algorithm (99), with all R-hat scores < 1.01. 95% credible intervals were calculated as the 95% highest posterior density interval (HPDI) using StatisticalRethinking.jl (https://github.com/StatisticalRethinkingJulia/StatisticalRethinking.jl). Subsequently, point estimates were obtained by computing the *maximum a posteriori* parameter estimates of the same model, using Optim.jl (100) in conjunction with Turing.jl. To compare the positive affect of copper on siderophore production amongst compost community isolates with the negative affect on SBW25, we computed the posterior distribution of the combined effect as β_1_ + β_2_ by element-wise addition of Markov chain samples. The same approach was used to infer the posterior distribution of the difference between community and SBW25 siderophore production with copper (α_2_ – α_1_) and without (α_2_ + β_2_ – (α_1_ + β_1_)).

#### The role of direct and indirect selection in favoring intermediate siderophore levels

To test for whether copper-adapted clones of SBW25 evolved greater resistance to copper than those evolved under unpolluted conditions, we used a LMM model on *m* with evolutionary background as fixed explanatory variable. To account for non-independency of observations, we fitted random intercepts for individual microcosms. However, the variation explained by the random intercepts approached zero, causing singularity issues when fitting the LMM using the ‘*lme4’* package. We therefore used the default setting of the nonlinear optimizer in the *‘glmmTMB’* function for parameter estimation. We next tested whether increased investment in siderophores conferred greater copper tolerance by including siderophore production in the evolutionary LMM described above.

We determined whether siderophores were selectively favored when iron was needed for growth by propagating a pyoverdine-producing and pyoverdine-deficient mutant strain of SBW25 in isolation and together at different inoculation densities under copper-polluted and unpolluted conditions. We first determined whether pyoverdines provide an *absolute* growth benefit to the wild-type producer under toxic copper conditions by comparing the Malthusian parameter (*m*) of individually grown strains using a GLM with density × strain × copper as explanatory variables, as well as all possible 2- and 3-way interactions. We next determined whether density differentially affected the performance of the pyoverdine producing and non-producing strains during direct competition, using a LMM with *m* as response variable and strain × frequency as explanatory variables and random intercepts fitted for each microcosm (*n* = 24) to account for non-independency of paired observations within microcosms. Finally, we were interested in how density affected the relative fitness of the pyoverdine-producing strain (𝑟_𝑝𝑟𝑜𝑑𝑢𝑐𝑒𝑟_ = 𝑚_𝑝𝑟𝑜𝑑𝑢𝑐𝑒𝑟_/𝑚_𝑛𝑜𝑛–𝑝𝑟𝑜𝑑𝑢𝑐𝑒𝑟_) (84) during direct competition with the mutant as a function of copper stress. We expected the pyoverdine-producing strain to be at a selective disadvantage under low growth conditions (i.e. when iron is not limiting growth). We tested this using a GLM with *r_producer_* as response variable and copper × density as explanatory variables, as well as their 2-way interaction.

Finally, we determined whether social exploitation could have played an additional role in selecting against high siderophore levels in response to copper stress. To this end, we competed the pyoverdine-producing wildtype strain SBW25 and a pyoverdine-deficient mutant at a low (1%) or high (99%) frequency in copper-polluted and unpolluted compost. We determined whether copper differentially affected the relative fitness of the pyoverdine-deficient mutant (𝑠_𝑚𝑢𝑡𝑎𝑛𝑡_ = 𝑚_𝑚𝑢𝑡𝑎𝑛𝑡_– 𝑚_*w*𝑖𝑙𝑑–𝑡𝑦𝑝𝑒_) (84) as a function of its initial frequency. Note that we used the selection rate (*s*) as a measure of relative fitness as some of the Malthusian growth parameters were negative (SI Appendix, Fig. S7). We expected the pyoverdine-deficient mutant to reap greater fitness benefits when rare and tested this hypothesis using a GLM with 𝑠_𝑚𝑢𝑡𝑎𝑛𝑡_ as response variable and initial frequency (low or high) × copper as fixed explanatory variables, as well as their interaction.

## Acknowledgments

We thank the three anonymous reviewers for their valuable comments and Daniel Padfield for advice on statistical analyses. This work was funded by AXA Research Fund, Biotechnology and Biological Sciences Research Council (BBSRC: BB/K00324) and Natural Environment Research Council (NERC: NE/V012347/1) grants awarded to AB and a UK Research Innovation (UKRI) Future Leader Fellowship (UKRI-FLF: MR/V022482/1) awarded to EH. SOB was funded by a ‘Bridging the Gaps’ award and PhD scholarship from the University of Exeter. AML was supported by FP7 Marie Skłodowska-Curie International Incoming Fellowship (IIF) (no. 331163).

## Author Contributions

EH, AML, SOB, DJH, AB conceived and designed the study. EH, AML, SOB, TM, JSP, FB collected the data. EH, AN and FB carried out data analyses.

EH, AB wrote the first draft of the manuscript, and all authors contributed to revisions.

## Competing Interest Statement

The authors declare no competing interests.

## Open Access

For the purpose of open access, the authors have applied a Creative Commons Attribution (CC BY) license to any Author Accepted Manuscript version arising from this submission’.

## Notes

https://zenodo.org/records/11639136

https://www.ebi.ac.uk/ena/data/search?query=PRJEB29924

### Competing Interest Statement

The authors have declared no competing interest.

### Summary of Updates

Following revision we have carried out additional experiments and analyses. This has altered the author list, and narrative of our work and has warranted a new title

## References

1. T. G. Barraclough, How do species interactions affect evolutionary dynamics across whole communities? Annu. Rev. Ecol. Evol. Syst. 46, 25–48 (2015).

2. J. Cairns, R. Jokela, L. Becks, V. Mustonen, T. Hiltunen, Repeatable ecological dynamics govern response of experimental community to antibiotic pulse perturbation. Nat. Ecol. Evol. 4, 1385–1394 (2020).

3. R. Evans et al., Eco-evolutionary dynamics set the tempo and trajectory of metabolic evolution in multispecies communities. Curr. Biol. 30, 4984–4988 (2020).

4. A. B. Chase, C. Weihe, J. B. Martiny, Adaptive differentiation and rapid evolution of a soil bacterium along a climate gradient. PNAS 118, e2101254118 (2021).

5. F. Fiegna, A. Moreno-Letelier, T. Bell, T. G. Barraclough, Evolution of species interactions determines microbial community productivity in new environments. ISME 9, 1235–1245 (2015).

6. N. G. Hairston Jr, S. P. Ellner, M. A. Geber, T. Yoshida, J. A. Fox, Rapid evolution and the convergence of ecological and evolutionary time. Ecol. Lett. 8, 1114–1127 (2005).

7. F. A. Gorter, M. Manhart, M. Ackermann, Understanding the evolution of interspecies interactions in microbial communities. Philos. Trans. R. Soc. Lond. B Biol. Sci. 375, 20190256 (2020).

8. J. B. Martiny et al., Investigating the eco-evolutionary response of microbiomes to environmental change. Ecol. Lett. 26, S81–S90 (2023).

9. M. Lässig, V. Mustonen, A. M. Walczak, Predicting evolution. Nat. Ecol. Evol. 1, 1–9 (2017).

10. N. Meroz, N. Tovi, Y. Sorokin, J. Friedman, Community composition of microbial microcosms follows simple assembly rules at evolutionary timescales. Nat. Commun. 12, 1–9 (2021).

11. Y. Yang, Emerging patterns of microbial functional traits. Trends Microbiol. 29, 874–882 (2021).

12. M. Cavaliere, S. Feng, O. S. Soyer, J. I. Jiménez, Cooperation in microbial communities and their biotechnological applications. Environ. Microbiol. 19, 2949–2963 (2017).

13. S. Shibasaki, S. Mitri, Controlling evolutionary dynamics to optimize microbial bioremediation. Evol. Appl. 13, 2460–2471 (2020).

14. A. N. Yadav et al., Beneficial microbiomes: biodiversity and potential biotechnological applications for sustainable agriculture and human health. J. appl. biol. 5, 4–7 (2017).

15. T. Scheuerl et al., Bacterial adaptation is constrained in complex communities. Nat. Commun. 11, 1–8 (2020).

16. M. Castledine, D. Padfield, A. Buckling, Experimental (co) evolution in a multi-species microbial community results in local maladaptation. Ecol. Lett. 23, 1673–1681 (2020).

17. D. Lawrence et al., Species interactions alter evolutionary responses to a novel environment. PLoS Biol. 10, e1001330 (2012).

18. A. Buckling, R. Craig Maclean, M. A. Brockhurst, N. Colegrave, The Beagle in a bottle. Nature 457, 824–829 (2009).

19. C. De Mazancourt, E. Johnson, T. Barraclough, Biodiversity inhibits species’ evolutionary responses to changing environments. Ecol. Lett. 11, 380–388 (2008).

20. P. Gómez, A. Buckling, Real-time microbial adaptive diversification in soil. Ecol. Lett. 16, 650–655 (2013).

21. M. A. Brockhurst, N. Colegrave, D. J. Hodgson, A. Buckling, Niche occupation limits adaptive radiation in experimental microcosms. PLoS One 2, e193 (2007).

22. R. S. Etienne, B. Haegeman, A conceptual and statistical framework for adaptive radiations with a key role for diversity dependence. Am. Nat. 180, E75–E89 (2012).

23. A. Jousset, N. Eisenhauer, M. Merker, N. Mouquet, S. Scheu, High functional diversity stimulates diversification in experimental microbial communities. Sci. Adv. 2, e1600124 (2016).

24. D. I. Bolnick et al., Why intraspecific trait variation matters in community ecology. TREE 26, 183–192 (2011).

25. R. Lanfear, H. Kokko, A. Eyre-Walker, Population size and the rate of evolution. TREE 29, 33–41 (2014).

26. J. Johansson, Evolutionary responses to environmental changes: how does competition affect adaptation? Evolution 62, 421–435 (2008).

27. R. Kummerli, K. T. Schiessl, K. McNeill, M. Ackermann, Habitat structure and the evolution of diffusible siderophores in bacteria. Ecol. Lett. 17, 1536–1655 (2014).

28. R. C. Hider, X. Kong, Chemistry and biology of siderophores. Nat. Prod. Rep. 27, 637–657 (2010).

29. C. Ratledge, L. G. Dover, Iron metabolism in pathogenic bacteria. Annu. Rev. Microbiol. 54, 881–941 (2000).

30. I. J. Schalk, M. Hannauer, A. Braud, New roles for bacterial siderophores in metal transport and tolerance. Environ. Microbiol. 13, 2844–2854 (2011).

31. A. Braud, V. Geoffroy, F. Hoegy, G. L. A. Mislin, I. J. Schalk, Presence of the siderophores pyoverdine and pyochelin in the extracellular medium reduces toxic metal accumulation in *Pseudomonas auruginosa* and increases bacterial metal tolerance. Environ. Microbiol. Rep. 2, 419–425 (2010).

32. S. O’Brien et al., No effect of intraspecific relatedness on public goods cooperation in a complex community. Evolution 72, 1165–1173 (2018).

33. E. Hesse et al., Ecological selection of siderophore-producing microbial taxa in response to heavy metal contamination. Ecol. Lett. 21, 117–127 (2018).

34. E. Hesse et al., Stress causes interspecific facilitation within a compost community. Ecol. Lett. 24, 2169–2177 (2021).

35. J. Kramer, Ö. Özkaya, R. Kümmerli, Bacterial siderophores in community and host interactions. Nat. Rev. Microbiol., 1–12 (2020).

36. E. Hesse et al., Anthropogenic remediation of heavy metals selects against natural microbial remediation. Proc B 286, 20190804 (2019).

37. P. B. Rainey, M. J. Bailey, Physical and genetic map of the Pseudomonas fluorescens SBW25 chromosome. Molecular Microbiology 19, 521–533 (1996).

38. A. M. Lujan, P. Gomez, A. Buckling, Siderophore cooperation of the bacterium *Pseudomonas fluorescens* in soil. Biol. Lett. 11, 20140934 (2015).

39. C. D. Moon et al., Genomic, genetic and structural analysis of pyoverdine-mediated iron acquisition in the plant growth-promoting bacterium *Pseudomonas fluorescens* SBW25. BMC Microbiol. 8, 7 (2008).

40. N. Jiricny et al., Fitness correlates with the extent of cheating in a bacterium. J. Evol. Biol. 23, 738–747 (2010).

41. A. S. Griffin, S. A. West, A. Buckling, Cooperation and competition in pathogenic bacteria. Nature 430, 1024–1027 (2004).

42. S. Pollak et al., Public good exploitation in natural bacterioplankton communities. Sci. Adv. 7, eabi4717 (2021).

43. J. Kramer, Ö. Özkaya, R. Kümmerli, Bacterial siderophores in community and host interactions. Nat. Rev. Microbiol. 18, 152–163 (2020).

44. J. Galet et al., *Pseudomonas fluorescens* pirates both ferrioxamine and ferricoelichelin siderophores from *Streptomyces ambofaciens*. Appl. Environ. Microbiol. 81, 3132–3141 (2015).

45. P. Cornelis, S. Matthijs, Diversity of siderophore-mediated iron uptake systems in fluorescent pseudomonads: not only pyoverdines. Environ. Microbiol. 4, 787–798 (2002).

46. R. Niehus, A. Picot, N. M. Oliveira, S. Mitri, K. R. Foster, The evolution of siderophore production as a competitive trait. Evolution 71, 1443–1455 (2017).

47. A. R. Figueiredo, Ö. Özkaya, R. Kümmerli, J. Kramer, Siderophores drive invasion dynamics in bacterial communities through their dual role as public good versus public bad. Ecol. Lett. 25, 138–150 (2022).

48. J. D. Palmer, K. R. Foster, Bacterial species rarely work together. Science 376, 581–582 (2022).

49. J. J. Morris, R. E. Lenski, E. R. Zinser, The Black Queen hypothesis: evolution of dependencies through adaptive gene loss. MBio 3, 2 e00036–00012 (2012).

50. W. D. Hamilton, The genetical evolution of social baviour. I. J. Theor. Biol. 7, 1–16 (1964).

51. L. J. Belcher, A. E. Dewar, M. Ghoul, S. A. West, Kin selection for cooperation in natural bacterial populations. PNAS 119, e2119070119 (2022).

52. M. Garcia-Garcera, E. P. Rocha, Community diversity and habitat structure shape the repertoire of extracellular proteins in bacteria. Nat. Commun. 11, 1–11 (2020).

53. O. X. Cordero, L.-A. Ventouras, E. F. DeLong, M. F. Polz, Public good dynamics drive evolution of iron acquisition strategies in natural bacterioplankton populations. PNAS 109, 20059–20064 (2012).

54. S. A. West, A. S. Griffin, A. Gardner, S. P. Diggle, Social evolution theory for microorganisms. Nat. Rev. Microbiol. 4, 597–608 (2006).

55. S. A. West, S. P. Diggle, A. Buckling, A. Gardner, A. S. Griffin, The social lives of microbes. Annu. Rev. Ecol. Syst. 38, 53–77 (2007).

56. B. J. Crespi, The evolution of social behavior in microorganisms. TREE 16, 178–183 (2001).

57. J. Gore, H. Youk, A. Van Oudenaarden, Snowdrift game dynamics and facultative cheating in yeast. Nature 459, 253–256 (2009).

58. G. Hardin, The tragedy of the commons. Science 162, 1243–1248 (1968).

59. R. Axelrod, W. D. Hamilton, The evolution of cooperation. Science 211, 1390–1396 (1981).

60. J. L. Sachs, U. G. Mueller, T. P. Wilcox, J. J. Bull, The evolution of cooperation. Q. Rev. Biol. 79, 135–160 (2004).

61. H. H. Lee, M. N. Molla, C. R. Cantor, J. J. Collins, Bacterial charity work leads to population-wide resistance. Nature 467, 82–85 (2010).

62. I. Frost et al., Cooperation, competition and antibiotic resistance in bacterial colonies. ISME 12, 1582–1593 (2018).

63. C. D. Nadell, J. B. Xavier, K. R. Foster, The sociobiology of biofilms. FEMS Microbiol. Rev. 33, 206–224 (2008).

64. K. P. Rumbaugh et al., Quorum sensing and the social evolution of bacterial virulence. Curr. Biol. 19, 341–345 (2009).

65. I. Eleftherianos et al., An antibiotic produced by an insect-pathogenic bacterium suppresses host defenses through phenoloxidase inhibition. PNAS 104, 2419–2424 (2007).

66. B. Raymond, S. A. West, A. S. Griffin, M. B. Bonsall, The dynamics of cooperative bacterial virulence in the field. Science 337, 85–88 (2012).

67. T. Köhler, G. G. Perron, A. Buckling, C. Van Delden, Quorum sensing inhibition selects for virulence and cooperation in *Pseudomonas aeruginosa*. PLoS Pathog. 6, e1000883 (2010).

68. P. Gómez et al., Local adaptation of a bacterium is as important as its presence in structuring a natural microbial community. Nat. Commun. 7, 12453 (2016).

69. R. Kummerli, A. S. Griffin, S. A. West, A. Buckling, F. Harrison, Viscous medium promotes cooperation in the pathogenic bacterium *Pseudomonas aeruginosa*. Proc B 276, 3531–3538 (2009).

70. E. Butaitė, M. Baumgartner, S. Wyder, R. Kümmerli, Siderophore cheating and cheating resistance shape competition for iron in soil and freshwater Pseudomonas communities. Nat. Commun. 8, 414 (2017).

71. A. Ross-Gillespie, A. Gardner, S. A. West, A. S. Griffin, Frequency dependence and cooperation: theory and a test with bacteria. Am. Nat. 170, 331–342 (2007).

72. H. Heuer, K. Smalla, Plasmids foster diversification and adaptation of bacterial populations in soil. FEMS Microbiol. Rev. 36, 1083–1104 (2012).

73. T. Robinson, P. Smith, E. R. Alberts, M. Colussi-Pelaez, M. Schuster, Cooperation and cheating through a secreted aminopeptidase in the *Pseudomonas aeruginosa* RpoS response. Mbio 11, 10.1128/mbio.03090-03019 (2020).

74. A. Dragoš et al., Evolution of exploitative interactions during diversification in Bacillus subtilis biofilms. FEMS Microbiol. Ecol. 94, fix155 (2018).

75. S. O’Brien, D. J. Hodgson, A. Buckling, Social evolution of toxic metal bioremediation in *Pseudomonas aeruginosa*. Proc B 281, 20140858 (2014).

76. A. K. Hottes et al., Bacterial adaptation through loss of function. PLoS genetics 9, e1003617 (2013).

77. T. Dimitriu et al., Genetic information transfer promotes cooperation in bacteria. PNAS 111, 11103–11108 (2014).

78. D. J. Rankin, E. P. Rocha, S. P. Brown, What traits are carried on mobile genetic elements, and why? Heredity (Edinb.) 106, 1–10 (2011).

79. A. E. Dewar, L. J. Belcher, T. W. Scott, S. A. West, Genes for cooperation are not more likely to be carried by plasmids. Proc B 291, 20232549 (2024).

80. T. Nogueira et al., Horizontal gene transfer of the secretome drives the evolution of bacterial cooperation and virulence. Curr. Biol. 19, 1683–1691 (2009).

81. O. X. Cordero, P. Hogeweg, The impact of long-distance horizontal gene transfer on prokaryotic genome size. Proceedings of the National Academy of Sciences 106, 21748–21753 (2009).

82. X. Cheng, I. de Bruijn, M. van der Voort, J. E. Loper, J. M. Raaijmakers, The Gac regulon of *Pseudomonas fluorescen*s SBW25. Environ. Microbiol. Rep. 5, 608–619 (2013).

83. B. Schwyn, J. B. Neilands, Universal chemical assay for the detection and determination of siderophores. Anal. Biochem. 160, 47–56 (1987).

84. R. E. Lenski, M. R. Rose, S. C. Simpson, S. C. Tadler, Long-term experimental evolution in *Escherichia coli*. I. Adaptation and divergence during 2000 generations. Am. Nat. 138, 1315–1341 (1991).

85. B. J. Callahan et al., DADA2: high-resolution sample inference from Illumina amplicon data. Nat. Methods 13, 581–583 (2016).

86. M. Kearse et al., Geneious Basic: an integrated and extendable desktop software platform for the organization and analysis of sequence data. Bioinformatics 28, 1647–1649 (2012).

87. P. D. Schloss et al., Introducing Mothur: open-source, platform-independent, community-supported software for describing and comparing microbial communities. Appl. Environ. Microbiol. 75, 7537–7541 (2009).

88. R. V. Lenth, Least-squares means: the R package lsmeans. J. Stat. Softw. 69, 1–33 (2016).

89. F. Hartig (2020) DHARMa: Residual diagnostics for hierachical (multi-level / mixed) regression models. R package version 0.3.3.0.

90. H. Wickham, ggplot2: Elegant Graphics for Data Analysis (Springer Verlag, New York, 2016).

91. D. Bates, M. Maechler, B. Bolker, S. Walker, Fitting Linear Mixed-Effects Models Using lme4. J. Stat. Softw. 67, 1–48 (2015).

92. J. Oksanen et al. (2010) vegan: Community Ecology Package. R package version 2.5–4.

93. M. I. Love, W. Huber, S. Anders, Moderated estimation of fold change and dispersion for RNA-seq data with DESeq2. Genome Biol. 15, 550 (2014).

94. M. E. Brooks et al., glmmTMB balances speed and flexibility among packages for zero-inflated generalized linear mixed modelling. The R Journal 9, 378–400 (2017).

95. M. Torchiano (2016) Effsize - A package for efficient effect size computation

96. A. Vehtari, A. Gelman, J. Gabry, Practical Bayesian model evaluation using leave-one-out cross-validation and WAIC. Stat. Comput. 27, 1413–1432 (2017).

97. H. Ge, K. Xu, Z. Ghahramani (2018) Turing: a language for flexible probabilistic inference. in International conference on artificial intelligence and statistics (PMLR), pp 1682–1690.

98. J. Bezanson, A. Edelman, S. Karpinski, V. B. Shah, Julia: A fresh approach to numerical computing. SIAM review 59, 65–98 (2017).

99. M. D. Hoffman, A. Gelman, The No-U-Turn sampler: adaptively setting path lengths in Hamiltonian Monte Carlo. J. Mach. Learn. Res. 15, 1593–1623 (2014).

100. P. K. Mogensen, A. N. Riseth, Optim: A mathematical optimization package for Julia. JOSS 3 (2018).

