## Supplementary Information for "Parallel ecological and evolutionary responses to selection in a natural bacterial community"

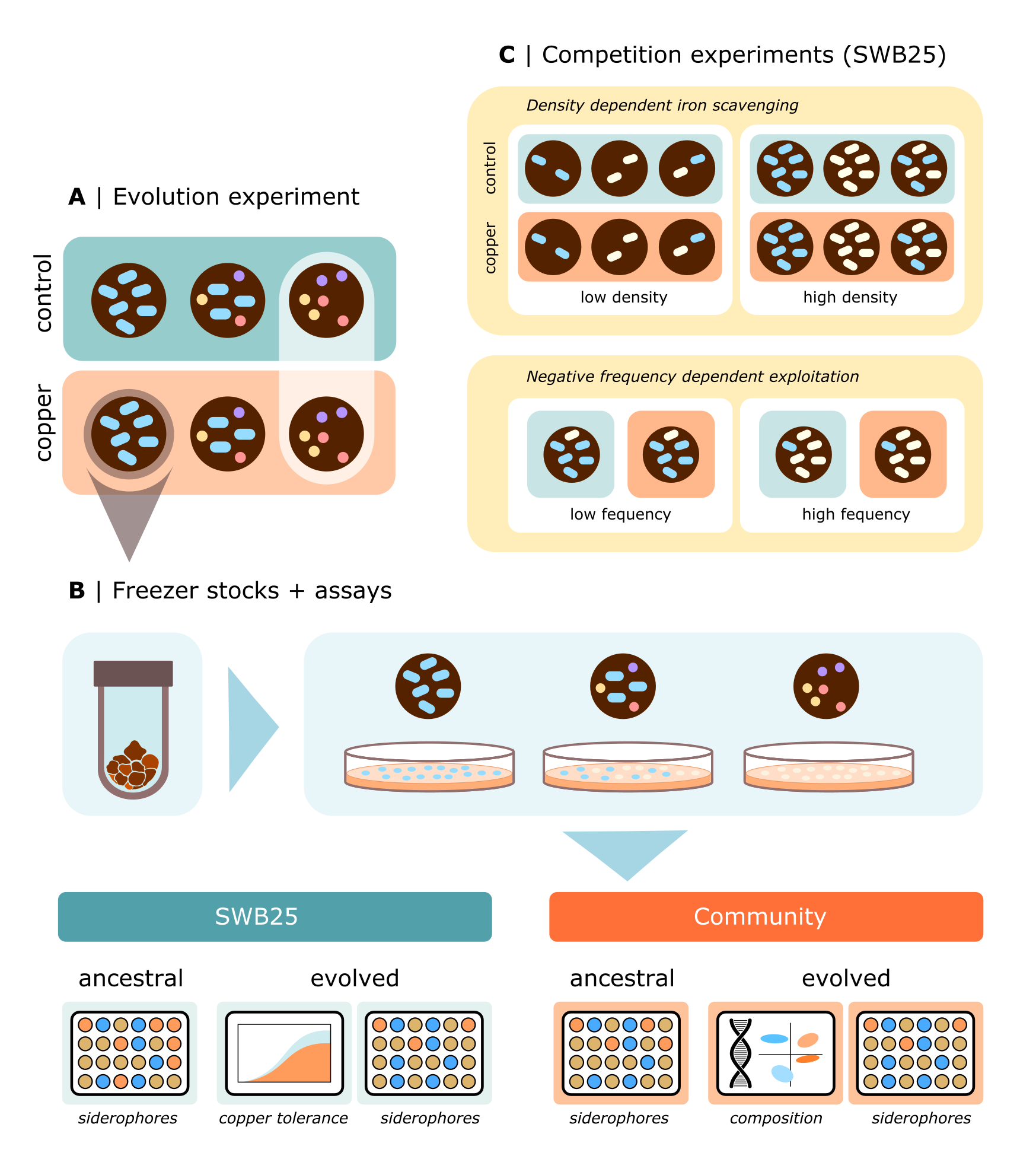

**Figure S1. Experimental design of the selection experiment and competition assays.** (**A**) We seeded twelve replicate microcosms with an overnight culture of SBW25 in sterile compost in the absence (blue ovals, left) or presence of the compost community (blue ovals + colored circles, middle). At the same time, we established twelve replicate compost communities by inoculating a soil wash (colored circles, right), keeping total inoculation density constant across treatments (~5 × 10^7^ cells). Note that the latter results (highlighted) have been published previously (32); here, we present the combined data set. After 24 hrs of incubation at 26°C, half of the replicate compost microcosms (copper, *n* = 6 per unique SBW25–community combination) received a toxic dose of copper (2 mL 0.25M CuSO_4_), and the remainder (unpolluted control, *n* = 6) an equal volume of sterile *dd*H2O. Compost communities were the incubated for 6 weeks. (**B**) We sampled all compost microcosms just before (ancestral) and 6 weeks after copper amendment (evolved) by mixing 1 gr of compost with 6 mL of sterile M9 solution, after which supernatants were frozen. Serial-diluted freezer stocks (*n* = 36) were plated on KB agar containing X-gal to distinguish SBW25 (blue colonies) from other community members (non-blue colonies). For each replicate, we randomly picked 24 colonies of either SWB25 and/or the community (i.e. 48 colonies in the SBW25 + community treatment) for phenotypic/genotypic assays, where possible (yielding 1049 clones in total). All colonies were individually grown in KB broth, and their siderophore production was assayed using colorific CAS assays. For isolated SBW25 clones that had evolved in the *absence* of the community, we also carried out copper tolerance assays (*n* =277. Note that for some populations we isolated < 24 clones). To determine whether siderophore changes in the community were due to ecological species sorting, we sequenced the 16S rRNA gene of all *evolved* compost isolates that had been assayed for siderophore production (*n* = 24 per replicate). (**C**) To test whether copper-imposed selection against high siderophore levels in SBW25 resulted from reduced competition for non-soluble iron, we carried out competition assays. In particular, we manipulated population growth rates by inoculating monocultures and co-cultures of wild-type LacZ marked SBW25 (blue) and an isogenic pyoverdine-deficient mutant (white) in sterile compost at very low density (allowing for faster growth) and high density (allowing for slower growth) (top panel) in copper-polluted and control compost. We next determined the role of social exploitation in driving copper-imposed reductions in siderophore production in SBW25 by carrying out another set of competition assays. In particular, we co-inoculated the wild-type pyoverdine-producing strain (blue) and pyoverdine-deficient mutant (white) at low or high frequency in copper-polluted (red highlight) and control (blue highlight) compost. We expected the pyoverdine-deficient strain to be at a selective advantage when initially rare. In both competition assays, we determined the Malthusian growth parameter of each strain by plating out serial-diluted soil washes on KB agar supplemented with X-gal, which resulted in blue colonies of the wild-type strain. Soil washes were collected after 7 days of growth, using the approach described in panel **B**.

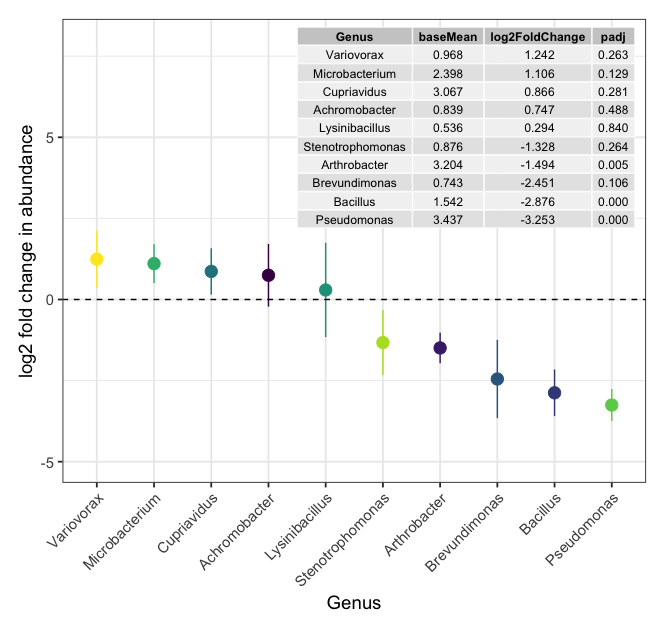

**Figure S2. The effect of SBW25 on the abundance of bacterial taxa in the compost community.** The log_2_-fold change in total abundance of the ten most common genera when comparing communities that had evolved in absence versus presence of SBW25. Of the taxa tested, three significantly differed in terms of total abundance between community-only versus community + SWB25 treatments: *Pseudomonas, Bacillus* and *Arthrobacter* were all negatively affected by the presence of SBW25 (see embedded table for Wald test and *P*-values for individual taxa, corrected for multiple testing using the ‘fdr’ method). Points represent estimated log_2-_fold change in total abundance of each taxon in response to evolving in the presence of SBW25 (averaged across copper regimes) ± standard error bars.

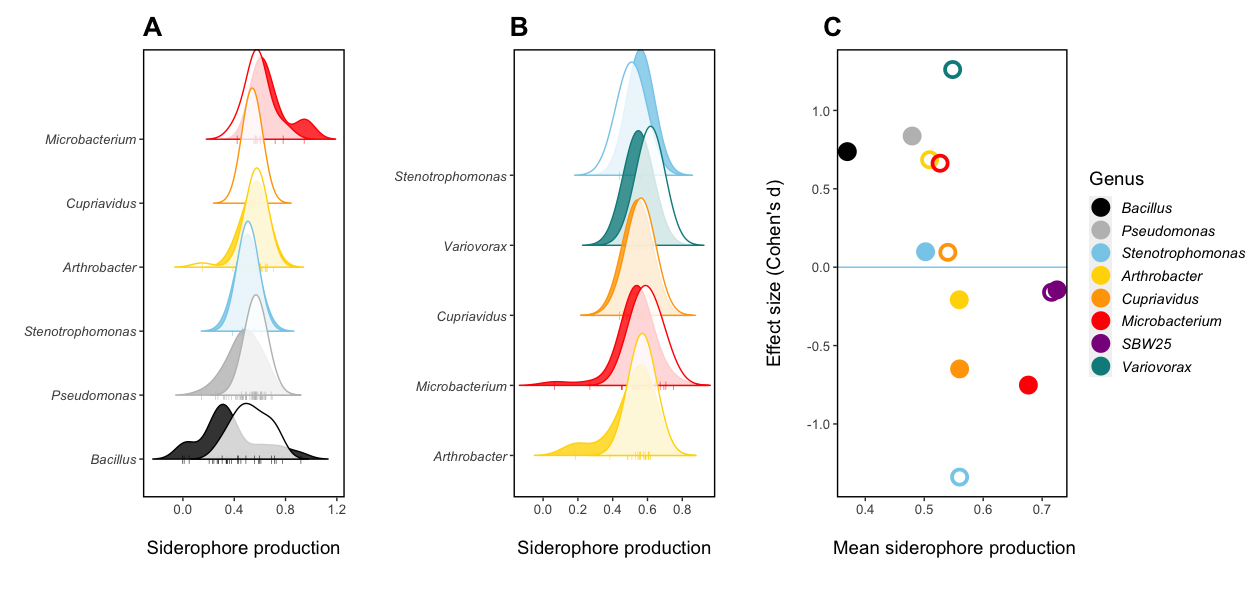

**Figure S3 | Copper pollution selects against siderophore extremes.** Panels **(A-B)** show the distribution of siderophore production in seven common bacterial compost taxa that had evolved in compost microcosms in the absence **(A)** or presence **(B)** of SBW25. Dark and light-shaded distributions depict unpolluted and copper-polluted communities, respectively. Panel **(C)** shows the negative correlation between effect size (Cohen’s *d* comparing mean siderophore production in copper-polluted versus unpolluted compost for seven compost taxa as well as SBW25) and mean ‘baseline’ siderophore production in unpolluted control compost. Open and closed symbols are estimates of Cohen’s *d* for taxa isolated from compost communities that had evolved for six weeks in the presence or absence of SWB25, respectively.

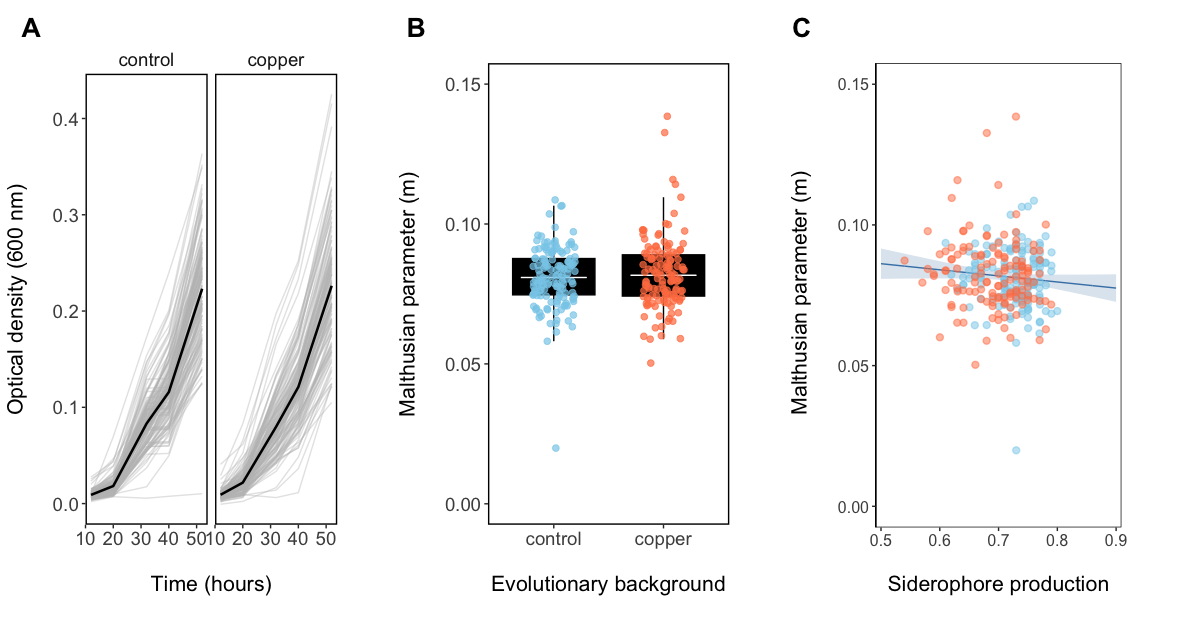

**Figure S4 | Selection for reduced siderophore production is not associated with direct fitness costs.** Panel (**A**) shows the effect of toxic copper (6.17 mM CuSO_4_) on temporal changes in the optical density (OD_600_) of individual SBW25 clones (grey lines) that had adapted to unpolluted control (*n* = 144 clones) or copper-polluted (*n* = 133 clones) conditions in the absence of the community. Bold black lines depict treatment-specific means. Panel (**B**) shows the Malthusian growth parameter (*m*) for unpolluted control (blue) and copper-adapted (red) clones in toxic copper broth. Boxes depict the upper and lower quartiles of raw data with the center line showing the median and whiskers providing a measure of 1.5x interquartile range. Note that we initially fitted replicate-specific random intercepts to accounts for non-independency of observations. However, the variance explained by the random effect approached zero; data for individual clones (colored points) are pooled across replicates for each copper background. (**C**) Plot showing the lack of relationship between copper resistance and siderophore production across different copper backgrounds (blue = unpolluted and red = copper-adapted clones). Line and blue shaded area depict the fitted linear relationship ± 95% confidence intervals (*m* = 0.10[0.08, 0.11] – 0.02[-0.05, 0] × siderophores).

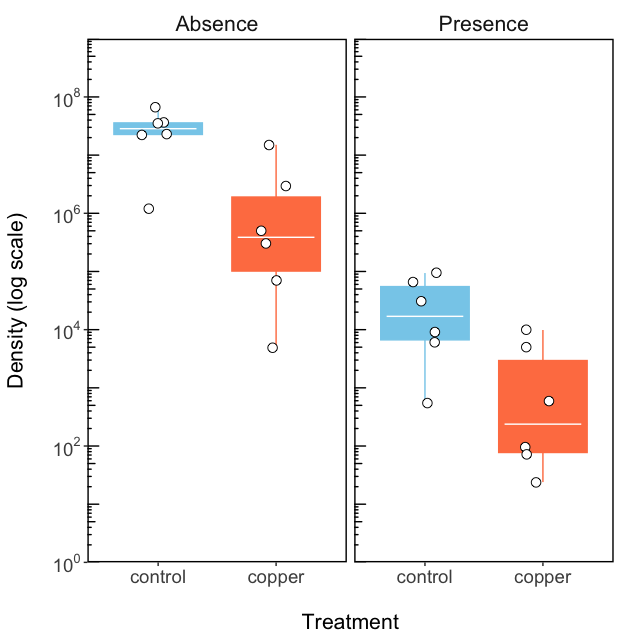

**Figure S5 | Copper and community presence reduce SBW25 density.** Six replicate populations were established in sterile compost in the absence or presence of the compost community and/or copper (unpolluted control *versus* copper-polluted compost), using a full factorial design. After 6 weeks of evolution, population densities were quantified by plating soil washes on LB agar supplemented with X-gal. Boxes depict the upper and lower quartiles of treatment-specific raw data (${log}_{10}$-cells gr^-1^ compost) with the center line showing the median and whiskers providing a measure of the 1.5x interquartile range. Points represent individual populations. Copper and community presence reduced final population density (*F*_1, 22_ = 63.53, *P* < 0.001 and *F*_1, 22_ = 17.88, *P* < 0.001 for main effects of community and copper). Letters denote significant Bonferroni-adjusted pairwise contrasts (z-ratio for SBW25 _control_ – SBW25 _copper_ = 4.23, SBW25_control_ – SBW25 + community _control_ = 7.97, SBW25 _control_ – SBW25 + community _copper_ = 8.63, SBW25 _copper_ – SBW25 + community _control_ = 2.65, SBW25 _copper_ – SBW25 + community _copper_ = 7.91 and SBW25 + community _control_ – SBW25 + community _copper_ = 4.23, all *P* < 0.05).

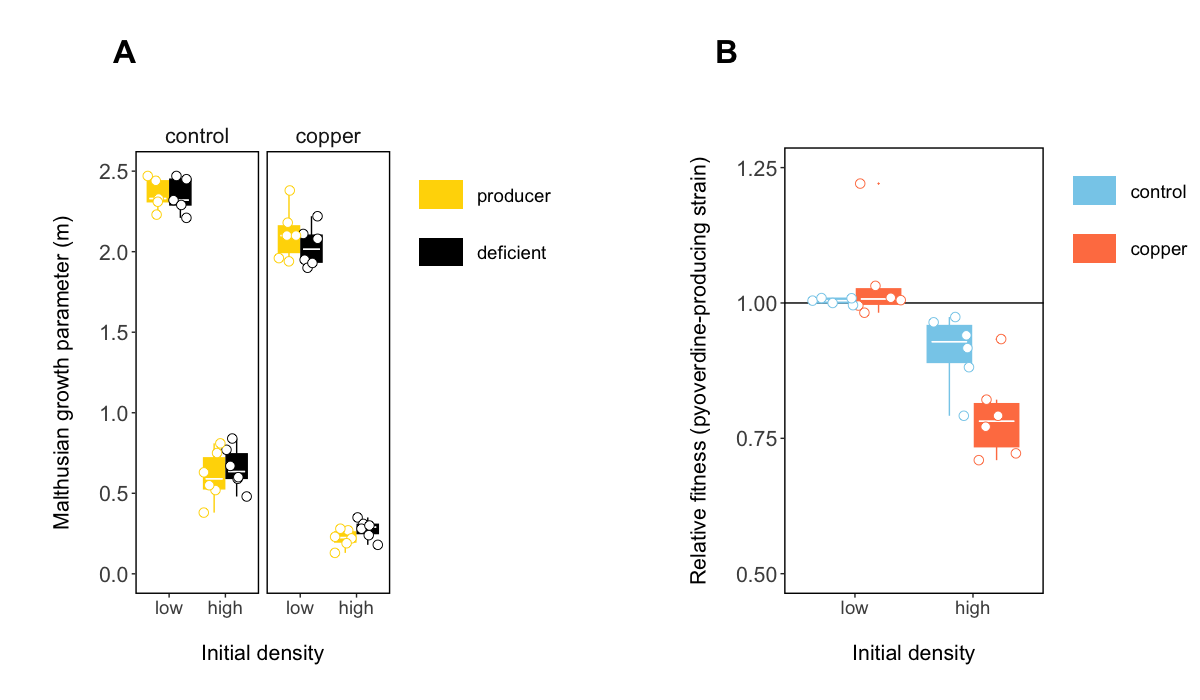

**Figure S6 | The effect of density on SBW25 in copper-polluted and unpolluted compost.** The pyoverdine-producing wildtype (gold) and pyoverdine-deficient mutant strain (black) were co-inoculated at 1:1 ratioin compost at low (10^2^ CFUs) or high (10^6^ CFUs) density. The effect of density on growth varied across strains (GLMM on *m*: density × strain: *χ_1,7_*= 6.57, *P* = 0.01), with the mutant growing better when growth was constrained ($\bar{m}_{producer}$ = 0.41 [0.35, 0.48] and $\bar{m}_{deficient}$= 0.47[0.40, 0.53], pairwise contrast: *t* ratio = -2.15, *P* <0.01), and the pyoverdine producer growing better under fast growth conditions ($\bar{m}_{producer}$= 2.24 [2.17, 2.31] and $\bar{m}_{deficient}$= 2.19 [2.12, 2.56], pairwise contrast: *t* ratio = 1.76, *P* = 0.09). Copper reduced population growth (*χ_1,6_*= 29.72, *P* < 0.001), irrespective of strain (copper × strain: *χ_1,8_*= 0.73, *P* = 0.39) and density differences (copper × density: *χ_1,7_*= 1.58, *P* = 0.22). Panel **(B)** demonstrates that the relative fitness of the pyoverdine-producing strain is greatest when competing with the non-producer under fast growth conditions ($r_{producer}$ > 1 when inoculation density was low), whilst the non-producer has a selective advantage under low growth conditions ($r_{producer}$ *<* 1 when inoculation density was high), in particular in copper-polluted conditions. Boxes depict the upper and lower quartiles of treatment-specific raw data with the center line showing the median and whiskers providing a measure of 1.5x interquartile range. Points show the raw data for each replicate population.

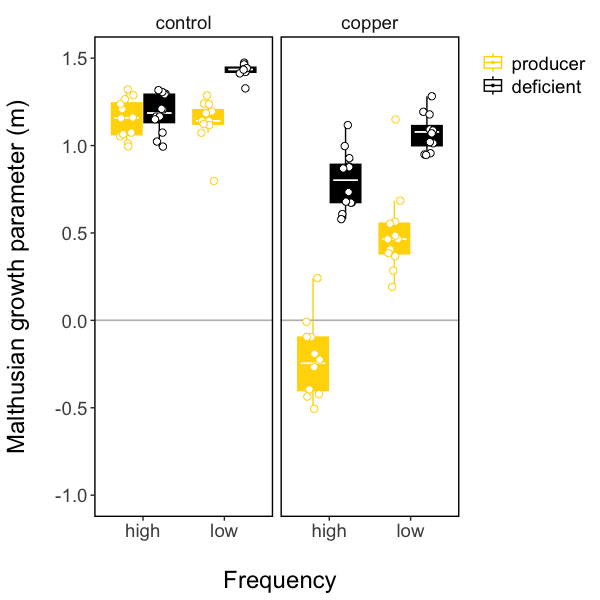

**Figure S7 | The effect of frequency on social exploitation in SBW25 in copper-polluted and unpolluted compost.** Boxplot depicting the Malthusian growth parameter (*m*) of the pyoverdine-deficient mutant (black) and pyoverdine-producing wild-type (gold) strains of SBW25 when inoculated together at different frequencies (low (1%) and high (99%) frequency) in copper-polluted and unpolluted control compost. The pyoverdine-deficient ‘cheat’ experienced a relative growth benefit, in particular when the producer was inoculated at a high frequency, and this effect was more pronounced under copper stress (LMM on *m*: effect of copper × strain × frequency on *m*: χ^2^_1_ = 20.25, *P* <0.001).

**Table S1. Change in the relative abundance of taxa in response to copper stress. Table shows mean across-treatment siderophore production, base mean abundance and log2 fold change for each taxon and accompanying *P*-values corrected for multiple testing using Wald tests (see methods).**

| **Genus** | **Siderophores** | **Base Mean** | **Log2 Fold Change** | **P_adj_** |
| --- | --- | --- | --- | --- |
| *Microbacterium* | 0.597 | 2.398 | -0.716 | 0.423 |
| *Variovorax* | 0.593 | 0.968 | -0.800 | 0.448 |
| *Cupriavidus* | 0.551 | 3.067 | 3.656 | **<0.001** |
| *Arthrobacter* | 0.548 | 3.204 | 0.482 | 0.448 |
| *Pseudomonas* | 0.527 | 3.437 | -0.849 | 0.307 |
| *Stenotrophomonas* | 0.519 | 0.876 | -0.443 | 0.710 |
| *Brevundimonas* | 0.516 | 0.743 | 1.623 | 0.304 |
| *Achromobacter* | 0.498 | 0.839 | -2.310 | 0**.028** |
| *Lysinibacillus* | 0.402 | 0.536 | -2.555 | **0.015** |
| *Bacillus* | 0.396 | 1.542 | -0.234 | 0.396 |
